# Inducible CRISPR/Cas9 allows for multiplexed and rapidly segregated single target genome editing in *Synechocystis* sp. PCC 6803

**DOI:** 10.1101/2022.03.02.482598

**Authors:** Ivana Cengic, Inés C. Cañadas, Nigel P. Minton, Elton P. Hudson

## Abstract

Establishing various synthetic biology tools is crucial for the development of cyanobacteria for biotechnology use, especially tools that allow for precise and markerless genome editing in a time-efficient manner. Here we describe a riboswitch-inducible CRISPR/Cas9 system, contained on one single replicative vector, for the model cyanobacteria *Synechocystis* sp. PCC 6803. A theophylline-responsive riboswitch allowed tight control of Cas9 expression, which enabled reliable transformation of the CRISPR/Cas9 vector into *Synechocystis*. Induction of the CRISPR/Cas9 mediated various types of genomic edits, specifically deletions and insertions of varying size. The editing efficiency varied depending on the target and intended edit; smaller edits overall performed better, reaching e.g. 100% for insertion of a FLAG-tag onto *rbcL*. Importantly, the single-vector CRISPR/Cas9 system described herein was also shown to mediate multiplexed editing of up to three targets in parallel in *Synechocystis*. All single-target and several double-target mutants were also fully segregated after the first round of induction, adding to the usefulness of this system. Further, a vector curing system that is separately induced by nickel and contained on the CRISPR/Cas9 vector itself, improved curing efficiencies by roughly 4-fold, enabling the final mutants to become truly markerless.

## Introduction

Cyanobacteria are photosynthetic prokaryotes that have gained growing interest as cell factories for sustainable production of various compounds.^1–3^ However, in order for such large-scale processes to become economically feasible, further study and significant engineering of these cyanobacterial hosts is required. While the toolset to engineer cyanobacteria is steadily growing,^4–6^ there is still need for reliable tools that allow for precise, markerless, rapid, and multiplexed editing in these polyploid organisms^7^ that are commonly time-consuming to engineer.

The CRISPR/Cas genome editing system is one such promising tool.^8–12^ A guide RNA (gRNA) is provided to guide the Cas endonuclease to a specific complementary target DNA site (the protospacer). This protospacer target must be next to a protospacer adjacent motif (PAM). Binding of the Cas-gRNA effector complex to the target DNA results in a double-stranded break (DSB) that is lethal for the cell unless mended. The most common method to mend DSBs in prokaryotes is by homology-directed repair (HDR), whereby a donor DNA is provided as a repair template. The Cas protein first popularized for widespread use was the class 2, type II Cas9 protein from *Streptococcus pyogenes*.^9,13^ However, besides being an efficient endonuclease it can also be cytotoxic when overexpressed, both on its own and particularly when co-expressed with a single guide RNA (sgRNA), causing low transformation efficiencies and thus lack of edited transformants.^14^ Some strategies to circumvent Cas9 toxicity are to identify and use alternative endonucleases, to engineer endogenous bacterial CRISPR-systems, or to decouple the transformation and editing steps by having inducible expression of Cas9.^14^

Several CRISPR/Cas systems have been described for cyanobacteria. In the two first studies, performed in *Synechococcus* UTEX 2973 and *Synechococcus elongatus* PCC 7942, transient expression of Cas9 supported editing, although cytotoxicity was a noted issue.^15,16^ In a later study, constitutive expression of the less toxic Cpf1 (also named Cas12a) endonuclease proved useful to mediate various types of edits (point mutation, knock-out, and knock-in) in UTEX 2973, *Synechocystis* sp. PCC 6803 (hereafter S6803), and *Anabaena sp*. PCC 7120.^17^ However, several passages on selective media were required to increase the percentage of edited and fully segregated colonies, especially in the highly polyploid S6803. This Cpf1-system was further evaluated in *Anabaena*, where it facilitated sequential and simultaneous deletion of two genes, and was supplemented with a counter-selection tool to cure edited cells of the Cpf1-plasmid.^18^ Only one example of an inducible CRISPR/Cas9 system has, to our knowledge, been described for cyanobacteria. There, *cas9* was integrated into the genome of the S6803 host and its expression was controlled by an aTc-inducible promoter.^19^ However, leaky Cas9 expression in un-induced cells caused low transformation efficiencies (∼10 CFU/µg DNA) for sgRNA-expressing plasmids. An alternative method using site-specific recombinases has also been used to make markerless genome edits in *Synechococcus* sp. PCC 7002 and S6803,^20^ however this method still leaves short recombination scars at the edit sites unlike CRISPR/Cas9 which supports scarless edits.

Development of tightly controlled CRISPR/Cas9 systems have until now been more successful in other types of bacteria.^14,21–23^ While many studies have used the traditional pairing of inducible promoter and transcriptional regulator, several recent studies have successfully employed riboswitches as regulatory elements for Cas9 expression.^24–26^ Riboswitches are small structured RNA-elements commonly found in the 5’-UTR of mRNAs.^27,28^ Their small size is especially useful when building CRISPR/Cas systems contained on single plasmids. The aptamer domain of a riboswitch is able to selectively bind to a specific ligand, triggering a conformational change that affects the expression platform domain and thus regulates the expression of the downstream gene.^27^ Often this regulation occurs at the transcriptional or translational level.

The riboswitches used to control CRISPR/Cas9 systems in bacteria have been based mainly on the synthetic theophylline-specific aptamer.^29^ This aptamer was used by Topp and coworkers to develop a set of six theophylline-inducible riboswitches (A-E + E*), widely applicable in different bacteria.^30^ These riboswitches regulate the expression of a downstream gene at the translational level by blocking access to the mRNA’s ribosome binding site (RBS) when no theophylline ligand is bound. When theophylline binds, the RBS is made available and translation can proceed. These theophylline riboswitches have already been evaluated in various cyanobacteria,^31–34^ and used in developing tools such as NOT-gates in e.g. S6803,^33^ inducible protein degradation systems in *S. elongatus* PCC 7942,^35,36^ and inducible CRISPR-interference systems in S6803 and *Anabaena*.^37,38^ In this study they were further used to design a tightly controlled CRISPR/Cas9 system for inducible genome editing in S6803.

## Results & Discussion

### Designing the riboswitch-based inducible CRISPR/Cas9 system for S6803

The goal of this study was to build a tightly controlled CRISPR/Cas9 system for S6803 where the expression of Cas9, and therefore genome editing, is controlled by a theophylline-inducible riboswitch. This system was built to be self-contained on a single plasmid based on the replicative pPMQAK1-T vector with an RSF1010-replicon.^39,40^

Of note is that the S6803 host used in this study is highly polyploid, with on average 20 genome copies observed during exponential growth.^7^ Any promising CRISPR/Cas9 system must therefore be able to edit all genomic copies of the intended target DNA to produce a fully segregated mutant. As the level of Cas9 required to achieve such editing in S6803 is unknown, three different riboswitches from the set developed by Topp *et al* ^30^ were tested to control its expression. The aim was to identify which would provide the correct balance of non-leaky Cas9 expression when un-induced, allowing high transformation efficiency of the CRISPR/Cas9 vector, and high enough expression when induced to allow for successful genome editing. The two riboswitches shown to have the least leaky expression in S6803, B and C,^33^ were selected. The more leaky variant E* was also selected,^33^ to include a riboswitch that supports a higher induced expression level if needed.

In a preliminary part of this study these riboswitches were combined with the *conII*-promoter (P_*conII*_) as has been done in several cyanobacteria studies.^32,33^ However, no useful system resulted from these efforts, mainly due to inefficiently induced CRISPR/Cas9 genome editing (data not shown). In a new attempt, the same riboswitches were instead combined with the *trc*-promoter (P_*trc*_) as described by Nakahira et al.;^31^ this also meant adding the P_*trc*_ 5’-UTR and a constant region upstream of the riboswitches. Note that the LacI-repressor was not expressed in any of the strains in this study so the inducibility of P_*trc*_ was only due to its combination with the riboswitches. To ensure that the theophylline concentrations used to induce the riboswitches wouldn’t be toxic, its toxicity towards S6803 was evaluated (Figure S1). At 0.5 mM theophylline there was no apparent growth defect, either in culture or on plates.

The effect of changing the promoter and 5’-UTR-region upstream of the riboswitches was studied with a Gfp-reporter (Figure S2). The P_*trc*_-variants supported higher expression levels and induction ratios than the P_*conII*_ ones. P_*trc*_-[B] and P_*trc*_*-*[E*] showed maximum induction ratios of 68- and 30-fold at 0.5 mM theophylline, respectively. The higher induction ratio for P_*trc*_-[B] compared to P_*trc*_-[E*] was due to less leaky expression, as P_*trc*_-[E*] allowed a 7-fold higher absolute expression level. In comparison, P_*trc*_-[C] underperformed due to leaky and weak expression and only reached a maximum 14-fold induction at 0.5 mM.

Two strategies to additionally lower the Cas9-levels in un-induced cells were explored. One was to add an *ssrA* protease degradation tag to the C-terminus of Cas9.^21^ The selected tag (LVA) has been estimated to reduce the steady-state level of a tagged protein by 95% in S6803.^39,41^ The other strategy was to mutate the -10-box in P_*trc*_ to emulate -10 boxes found in weaker promoters.^42^ The P_*trc*_ shares the same -10-box (TATAAT) as the strong BioBrick BBa_J23119 promoter, which belongs to a promoter library that has been evaluated in S6803.^40^ Promoters from this library that drive weaker expression and that only differ in their -10-box sequence were used to select the alternative -10-boxes. These selected ones came from promoters BBa_J23104 (TATTGT), BBa_J23116 (GACTAT), and BBa_J23117 (GATTGT). These were hereafter identified by their last three digits (104, 116, 117) and were estimated to weaken P_*trc*_ by roughly 65%, 90 %, 99%, respectively. These two weakening strategies were applied separately from each other to all P_*trc*_-riboswitch (B, C, or E*) pairs.

In total this created 15 different pPMQAK1-CRISPR/Cas9 base vectors to evaluate. All vectors were constructed in the same way for ease of use; the *cas9* expression cassette was followed by a *lacZ* sandwiched between two Golden Gate cloning BsaI sites (Figure 1a). The pieces specific to the target DNA, the sgRNA and donor DNA, were constructed with compatible BsaI sites (Figure 1a), enabling simultaneous ligation of both into the base vector to create the final target vector (Figure 1b).

**Figure 1.**
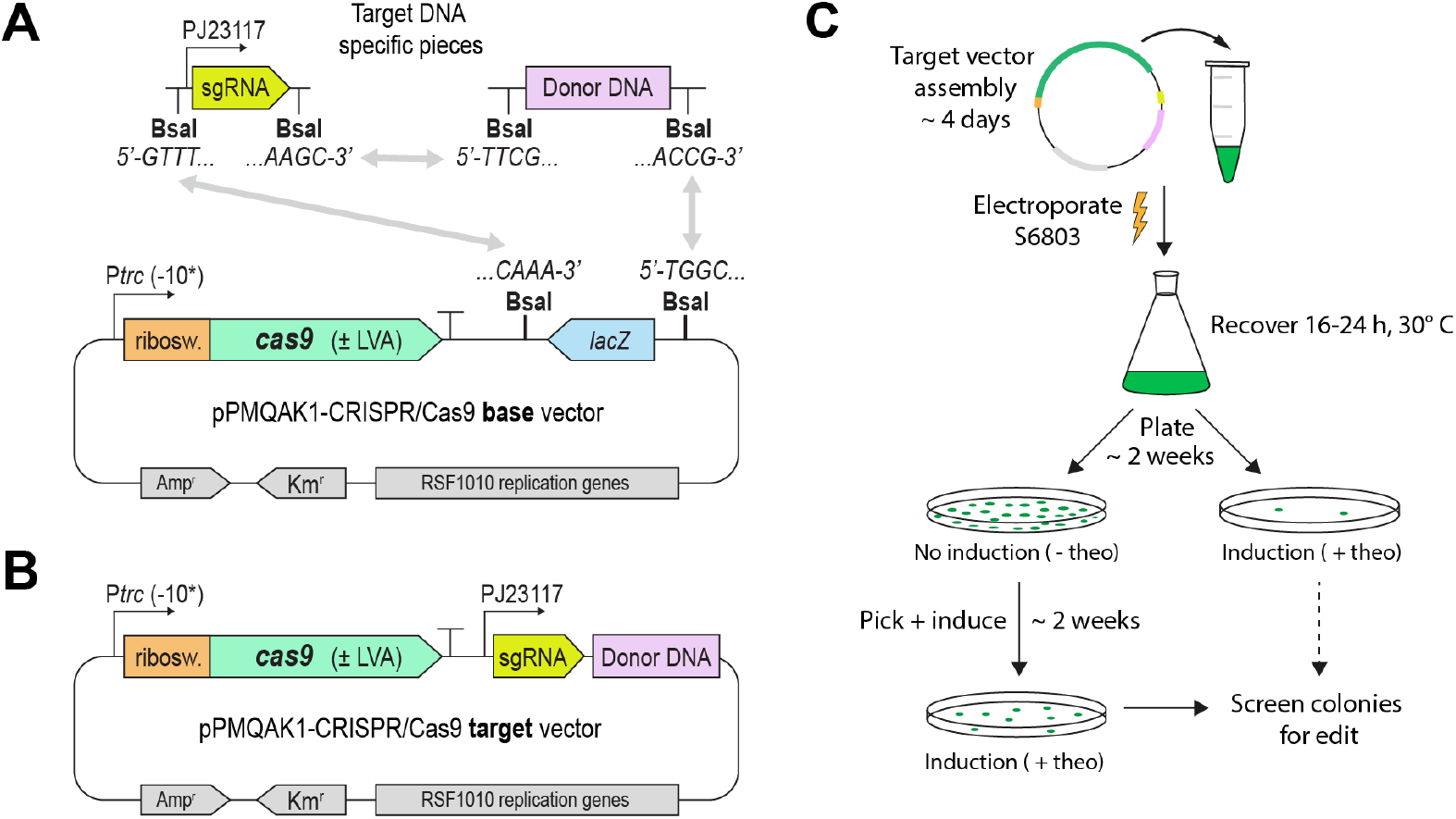
Overview of the pPMQAK1-CRISPR/Cas9 vector build and general workflow. (**A**) Schematic showing the pPMQAK1-CRISPR/Cas9 base vector and target DNA specific pieces, i.e. sgRNA and donor DNA, with the BsaI-overhangs required for one-step Golden Gate assembly. The vector variants differ in their expression of *cas9*: one of three riboswitches (“ribosw.”) mediate induction with theophylline, P_*trc*_ has -10-boxes with differing strengths (−10*), and some variants include an LVA-tagged Cas9 (±LVA). (**B**) Map of the final pPMQAK1-CRISPR/Cas9 target vector. (**C**) Workflow to transform a constructed pPMQAK1-CRISPR/Cas9 target vector into S6803 and induce CRISPR/Cas9-mediated editing. Approximate time for each step is indicated.

The promoter selected for sgRNA expression was the constitutive but weak BioBrick BBa_J23117.^40^ Weak sgRNA expression was deemed to complement the inducible CRISPR/Cas9 system better, since unnecessary overexpression could exacerbate the action of any leaky Cas9. To select the spacers used in the sgRNAs, the Cas9-specific NGG-PAM was used to search the target DNA area for suitable protospacers. The sgRNAs were designed to target as close to the desired edit site as possible, considering that Cas9 cuts 3-4 nucleotides upstream of the PAM.^12^ A short distance between the DSB and edit site is correlated with better editing efficiency.^43^ The sgRNAs were also constructed to target the template strand only; this allows for faster dislodging of the Cas9-sgRNA complex from the cut site, enabling improved access for the HDR-machinery.^44^

Depending on the desired genome edit, the donor DNA was altered accordingly. Generally, the donor DNA included 350 bp homology arms on either side of the edit site, and was designed to remove or silently mutate the PAM and proximal protospacer sequence to avoid recognition and continued cutting of the target site after editing.

The workflow (Figure 1c) used in this study took advantage of the decoupling of transformation and genome editing made possible by having inducible Cas9 expression. Plating of transformed and recovered S6803 on non-inducer plates resulted in many transformants for the pPMQAK1-CRISPR/Cas9 target vector. These transformants were subsequently plated on inducer-plates to undergo CRISPR/Cas9-mediated genome editing. Surviving colonies were screened for the desired edit. To control that the constructed pPMQAK1-CRISPR/Cas9 target vector was functional in mediating DSBs, i.e. had an efficient sgRNA, half of the transformed and recovered S6803 were plated directly on inducer-plates. If the DNA targeting was functional, these yielded no or very few surviving transformants due to the lethality of DSBs and low efficiency of HDR. If any transformants survived they were screened for the desired edit.

### Proof-of-principle test to identify promising CRISPR/Cas9 base vectors

To narrow down the set of the 15 designed pPMQAK1-CRISPR/Cas9 base vectors, a proof-of-principle test was done. The goal was to introduce a small 20 bp deletion into the *yfp*-gene of a S6803 Δ*slr1181*::P_*psbA2*_-Yfp-B0015-Sp^r^ strain (Figure 2a). An sgRNA targeting close to the middle of *yfp* was selected, while the desired 20 bp deletion included removal of the GGG-PAM and preceding three bases of the protospacer. The donor DNA contained homology arms surrounding the deletion site. All 15 resulting CRISPR/Cas9 *yfp*-target vectors were transformed into strain Δ*slr1181*::P_*psbA2*_-Yfp-B0015-Sp^r^ and treated according to the workflow in Figure 1c. A control vector where *cas9* lacked a promoter and start codon, i.e. resulting in no expression, was also included. This vector served as a control for the toxicity exhibited by any potentially leaky Cas9 expression in the tested vector variants. Its similarly large size (only 160 bp smaller) also worked as a control for the transformation efficiency for such large vectors (∼13 kbp) into S6803.

**Figure 2.**
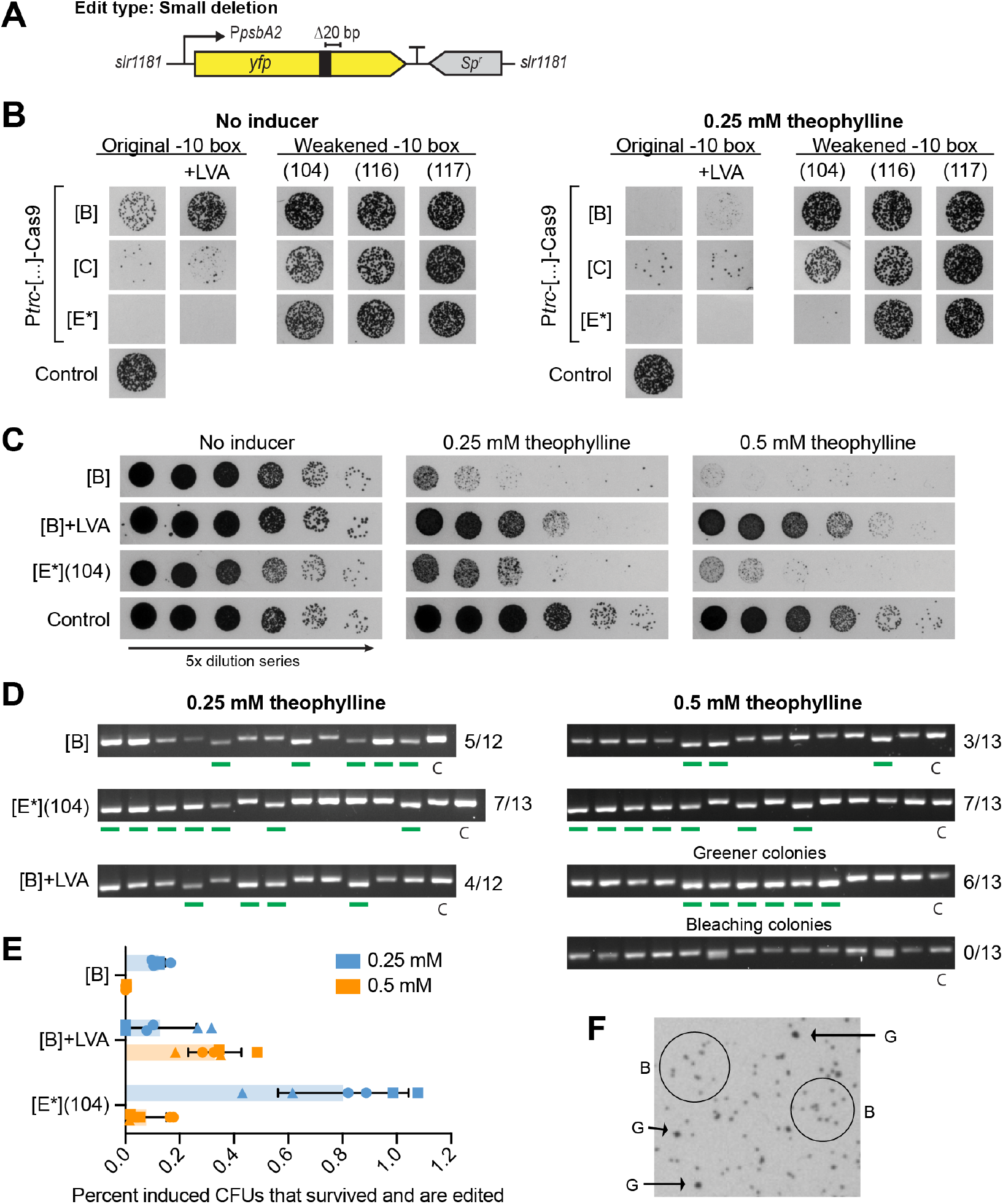
Evaluating the built pPMQAK1-CRISPR/Cas9 vectors in S6803 by performing a small deletion. (**A**) Schematic of the genomic *yfp*-target in S6803 Δ*slr1181*::P_*psbA2*_-Yfp-B0015-Sp^r^, showing approximate placement of the sgRNA target region (black box) and desired 20 bp deletion. (**B**) Transformation results after plating on selective plates without or with 0.25 mM theophylline inducer. The 15 *yfp*-targeting pPMQAK1-CRISPR/Cas9 vector variants differ in their *cas9*-expression strength, the P_*trc*_ is fused to riboswitch B, C, or E* and evaluated as is, in combination with a degradation tag (+LVA) on Cas9, or with a weakened P_*trc*_ by changing its original -10-box (strength: original>104>116>117). The control lacks *cas9* expression. (**C**) Induction spot assay results for [B], [B]+LVA, [E*](104), and the control. 5x dilution series were plated on regular BG11-plates or ones supplemented with 0.25 or 0.5 mM theophylline inducer. Done for biological triplicates in technical duplicates, representative data is shown. (**D**) Editing results for [B], [B]+LVA, [E*](104). A green line below a lane signals a fully edited (Δ20 bp) mutant. A control (“C”) shows how an unedited colony will appear. Fractions indicated the number of fully edited colonies out of the total number screened. Two representative colony phenotypes were screened for [B]+LVA induced on 0.5 mM theophylline. (**E**) Percentage of CFUs that survived and became edited, after being induced on 0.25 or 0.5 mM theophylline. Bars show the averages ± SD for biological triplicates (different symbol for each triplicate) done in technical duplicates. (**F**) Example of the two representative phenotypes found on inducer-plates, i.e. larger, greener colonies (indicated by arrows and marked “G”), and bleaching colonies (circled and marked “B”).

The desired result was a CRISPR/Cas9 target vector that resulted in many transformants under non-induced conditions, but was proven effective in inducing lethal DSBs when induced with theophylline. The transformation results (Figure 2b) clearly singled out constructs P_*trc*_-[B]-Cas9 (hereafter named [B]), P_*trc*_-[B]-Cas9+LVA (hereafter named [B]+LVA), and P_*trc*_(−10-box:104)-[E*]-Cas9 (hereafter named [E*](104)) as having these desirable traits. Based on the previous Gfp-reporter results (Figure S2), and what is known about the modifications introduced to weaken the expression or stability of Cas9 in these constructs, these systems were estimated to rank as follows in terms of resulting Cas9 amounts: [E*](104)>[B]>[B]+LVA. The control showed that 0.25 mM theophylline alone did not affect the transformation efficiency.

Colonies from the transformation plates (Figure S3a) were screened to check for leaky editing in the absence of any inducer, and if the few surviving transformants on the theophylline plates (Figure S3a-b) were indeed edited or had remained un-edited by somehow “escaping” the CRISPR/Cas9. While no leaky editing was observed for the un-induced transformants, a fraction of the surviving induced transformants were fully edited, with editing efficiencies of 25-87.5% depending on the construct (Figure S3c). This showcases the tight control of these CRISPR/Cas9 systems, and also the possibility of obtaining fully edited transformants directly after electroporation.

To further evaluate the CRISPR/Cas9-induction and *yfp* (Δ20 bp) editing efficiency of the promising [B], [B]+LVA, and [E*](104) constructs, triplicate transformants were induced on plates with theophylline. The promoterless-*cas9* vector was included as a control. In an attempt to titrate the Cas9 expression, induction on 0.25 mM and 0.5 mM theophylline was compared. The resulting representative spot assays (Figure 2c) expectedly showed widespread cell death due to induction of the CRISPR/Cas9 system. To determine the *yfp* (Δ20 bp) editing efficiency, surviving colonies that appeared healthy on the inducer-plates were picked and screened by colony-PCR (Figure 2d). While no construct reached 100% editing efficiency, all edited colonies were fully segregated after just one round of induction. In addition, the editing efficiencies for the two tested theophylline concentrations were comparable, showing that higher induction was not necessary. Overall, the stronger Cas9 expressing construct [E*](104) supported the best *yfp* (Δ20 bp) editing efficiency, reaching 54% for both tested inducer concentrations. To better judge the difference between the two tested theophylline concentrations and CRISPR/Cas9 constructs, the respective editing efficiency for the screened colonies was related to the percentage of CFUs that survived induction (see method section for details). This quantified the total percentage of induced CFUs that survived and became edited. The weakest [B]+LVA construct was found to benefit from increased Cas9 expression at 0.5 mM theophylline (Figure 2e). Meanwhile, induction with 0.25 mM outperformed 0.5 mM for constructs [B] and [E*](104) (Figure 2e), due to fewer colonies surviving on the higher concentration (Figure 2c). Taken together, the decision was made to use the lower 0.25 mM theophylline concentration for the rest of this study.

Among the colonies that survived CRISPR/Cas9-induction, two different phenotypes were observed. One type was greener and larger, while the second was smaller and slightly bleached (Figure 2f). This second phenotype was more prevalent for the weakest [B]+LVA construct and was responsible for the apparent higher survival of these cells on theophylline (Figure 2c). When these two colony types were compared in terms of *yfp* (Δ20 bp) editing for construct [B]+LVA, only the larger, greener ones exhibited any fully segregated edits (Figure 2d). Also, after prolonged induction on theophylline the smaller colonies were found to bleach entirely (Figure S3d), likely due to incomplete or absent editing and continued exposure to lethal Cas9-catalyzed DSBs. This phenotypic difference can therefore be useful when choosing which surviving colonies to screen. This strategy was applied throughout this continued study; only the larger, greener, and therefore healthier-looking colonies were assessed for genome editing.

### Exploring the edit types possible with this inducible CRISPR/Cas9 tool

The promising [B], [B]+LVA, and [E*](104) CRISPR/Cas9 vectors were further evaluated for their ability to perform other types of genomic edits in S6803.

To perform a large deletion, the entire P_*psbA2*_-Yfp-B0015-Sp^r^ cassette (Δ2240 bp) in the Δ*slr1181*::P_*psbA2*_-Yfp-B0015-Sp^r^ strain was targeted (Figure 3a). The same *yfp*-targeting sgRNA as used previously for the small 20 bp deletion was reused here. Although this sgRNA binds in the middle of *yfp* (Figure 3a), far from the edit site, such targeting has been shown to work for CRISPR/Cpf1 deletions in *Anabaena*.^18^ The donor DNA was changed to feature 350 bp homology arms on either side of the *slr1181* integration site. Transformation of the [B], [B]+LVA, and [E*](104) *yfp*-targeting vectors into S6803 resulted in many transformants (Figure S4). Induction of the CRISPR/Cas9 gave few surviving colonies (Figure 3b), the weakest [B]+LVA construct yielded only bleaching colonies (data not shown) and was not considered further for this target. For induced [B] and [E*](104) transformants, the editing efficiencies were 67% and 33%, respectively (Figure 3c). Despite the higher editing efficiency for construct [B], the total percentage of induced CFUs that survived and became edited was higher for the stronger construct [E*](104) (Figure 3d). One outlier for [E*](104) was especially successful.

**Figure 3.**
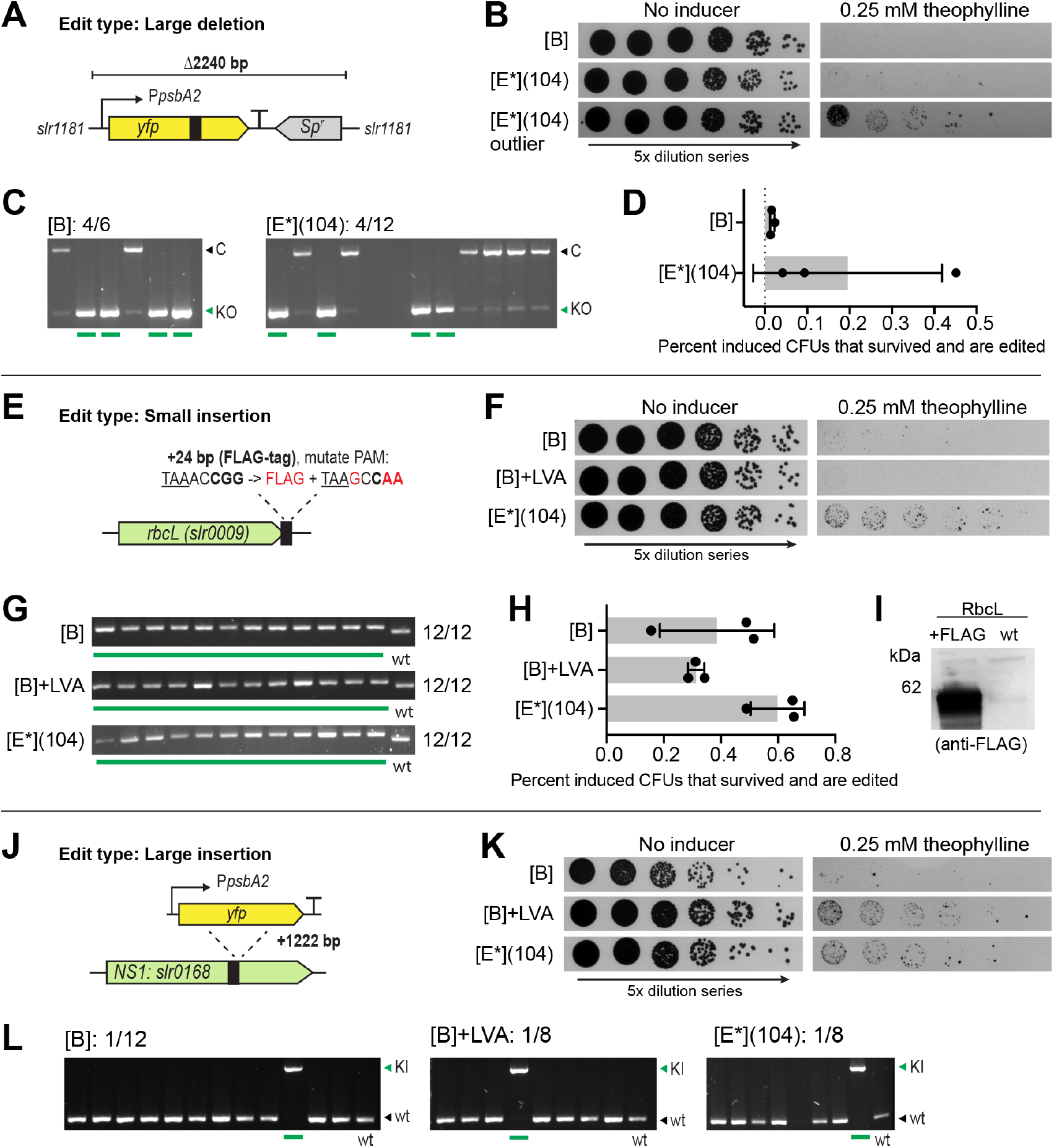
Testing various CRISPR/Cas9-edit types in S6803: (**A-D**) large deletion of a P_*psbA2*_-Yfp-B0015-Sp^r^ cassette, (**E-I**) small insertion of a FLAG-tag onto *rbcL*, (**J-L**) and large insertion of a P_*pbsA2*_-Yfp-B0015 cassette. (**A, E, J**) Schematics of the different targets in S6803, showing approximate placement of the sgRNA target region (black box) and intended edits. For (**E**) the TAA-stop codon is underlined, the CGG-PAM shown in bold. Mutations introduced in the edit are shown in red. (**B, F, K**) Induction spot assay results for specified CRISPR/Cas9 constructs. 5x dilution series were plated on plates without or with 0.25 mM theophylline. Done for biological triplicates, representative data is shown. (**C, G, L**) Editing results for specified CRISPR/Cas9 constructs. A green line below a lane signals a fully edited mutant. Fractions indicated the number of fully edited colonies out of the total number screened. For (**C**) “neg” and “KO” indicate the band-size for an un-edited and edited colony, respectively. For (**L**) “wt” and “KI” indicate the band-size for an un-edited and edited colony, respectively. (**D, H**) Percentage of induced CFUs that survived and became edited. Bars show the averages ± SD from biological triplicates, individual values are also shown. (**I**) Western blot of a fully edited mutant expressing RbcL-FLAG, compared to a wt control, 15 µg protein from the soluble fraction was loaded and probed using a anti-FLAG IgG.

Next a small genomic insertion was tested. The aim was to add a C-terminal FLAG-tag (24 bp) to the large subunit of Rubisco (*rbcL*) (Figure 3e). Many proteins of interest lack available antibodies, so the ability to introduce a markerless tag that does not disrupt the region surrounding the gene is a valuable tool. The position of this edit needed to be more specific than the previous deletions, as the FLAG-tag must be added in-frame in front of the *rbcL* stop codon (TAA). The protospacer options in this region were evaluated, and an sgRNA was designed to introduce the DSB two nucleotides from the desired edit site. As the CGG-PAM and parts of the proximal protospacer could not be deleted in this instance, they were instead mutated to stop the edited target from being recognized by the Cas9-sgRNA complex (Figure 3e). Transformation of the [B], [B]+LVA, and [E*](104) *rbcL*-target vectors into S6803 resulted in many transformants (Figure S5a). For constructs [B]+LVA and [E*](104), colonies that survived on the inducer-plates directly after transformation were already fully edited (Figure S5b). Inducing CRISPR/Cas9 in the transformants gave few surviving colonies (Figure 3f), but enough so the healthy colony phenotype could be found for all three tested vectors (Figure S5c). The editing efficiency was found to be 100% for all three *rbcL*-targeting constructs (Figure 3g). The higher editing efficiency supported for this edit, compared to the *yfp*-one previously, is likely due to a higher on-target efficiency of the *rbcL*-targeting sgRNA. Despite this, the total percentage of induced CFUs that survived and became edited was not much higher (Figure 3h). The soluble protein fraction from one of the edited colonies was subjected to immunoblotting using an anti-FLAG IgG. Here the product of the edited *rbcL*, i.e. the RbcL-FLAG, could be clearly detected (Figure 3i).

As a final test, an attempt was made to insert the whole P_*psbA2*_-Yfp-B0015 cassette into neutral site *slr0168* (Figure 3j). This cassette lacked any antibiotic marker, meaning that selection was done only by colonies surviving the Cas9-induced DSBs. The sgRNA was designed to target *slr0168*, and the donor DNA was supplied as three separate pieces when assembling the target vector. One piece for the whole P_*psbA2*_-Yfp-B0015-cassette, and one piece each for the upstream and downstream homologous regions (350 bp each). The donor DNA was also designed to remove the TGG-PAM and preceding three bases of the protospacer. Transformation of the [B], [B]+LVA, and [E*](104) target vectors into S6803 resulted in transformants (Figure S6), but fewer than seen for other target vectors previously. Induction of CRISPR/Cas9 gave many surviving colonies that exhibited the healthier phenotype (Figure 3k). However, editing efficiencies amongst the screened colonies was low (Figure 3l), as only a single fully edited colony (+1222 bp) was found per construct. It has been shown in *E. coli* that CRISPR/Cas9 editing efficiency is negatively correlated with an increase in insertion length,^45^ likely explaining the lower efficiency seen for this Yfp-cassette insertion. This efficiency could possibly be improved by extending the length of the homology arms in the donor DNA,^45^ however this was not explored here.

### Inducible multiplexed CRISPR/Cas9 editing in S6803

Multiplexed, simultaneous editing capabilities could greatly accelerate strain engineering in S6803. Considering the higher editing efficiencies of the smaller edits described above, these were selected for multiplexing tests. Such small edits can be used to modify promoter regions to alter gene expressions, or to introduce premature stop codons to knock-out genes.

As a proof-of-principle study the preliminary goal was to target four neutral sites (NS) in the S6803 genome and introduce small deletions (Δ30 bp) into all of them with a single pPMQAK1-CRISPR/Cas9 target vector. The NS were NS1 (*slr0168*), NS2 (*slr1181*), NS3 (*slr2030*-*2031*), and NS4 (*slr0397*); see Figure 4a for the respective sgRNA target regions. The donor DNAs had 350 bp homology arms; the 30 bp deletions included removal of the PAM-sites and preceding 3 bases of the protospacers.

**Figure 4.**
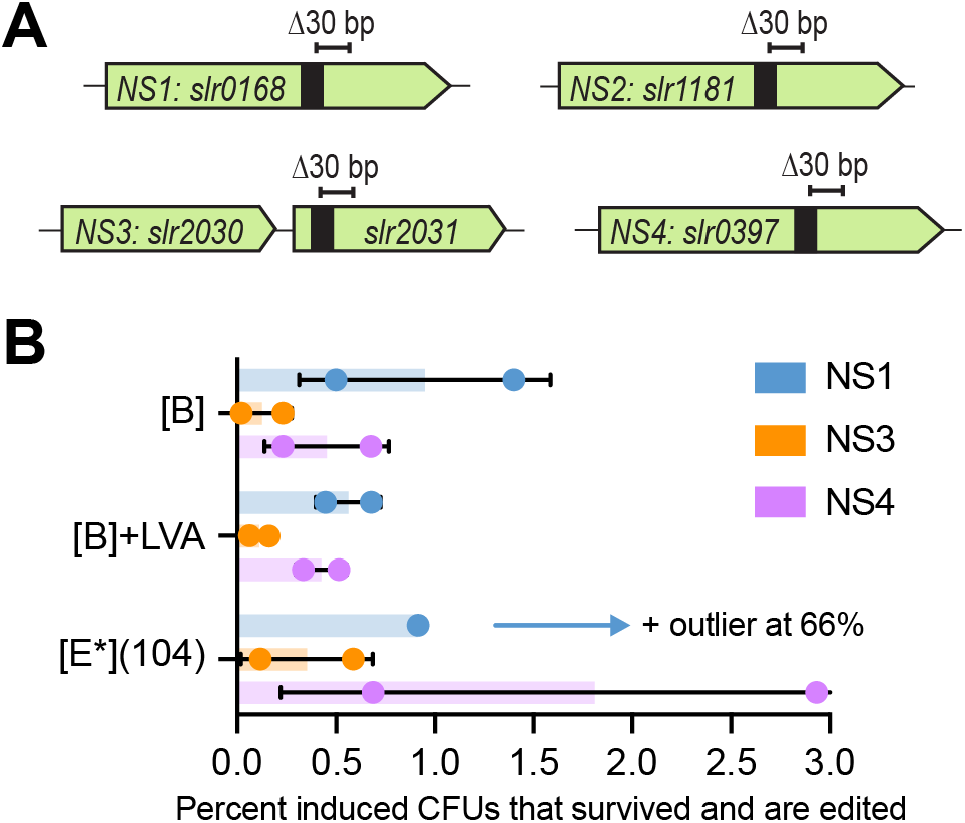
(**A**) Schematics of the S6803 neutral site (1-4) targets, showing approximate placement of the sgRNA target region (black box) and intended edit (Δ30 bp). (**B**) Percentage of induced CFUs that survived and became edited, for the individual editing of the NS1, NS3, and NS4 targets. Bars show the averages ± SD from biological duplicates, individual values are also shown. An outlier for [E*](104), target NS1, is not included in the graph but indicated by an arrow.

The sgRNA and donor DNA for each of the four targets was first tested individually with the [B], [B]+LVA, and [E*](104) base vectors. Many transformants were obtained for most constructs, with the exception of the NS1- or NS4-targeting [E*](104) constructs (Figure S7). Likely the sgRNAs designed for these two targets are highly efficient, exacerbating the effect of any leaky Cas9 expression from the stronger [E*](104) construct. Induction of CRISPR/Cas9 in these single target transformants gave varying results depending on the target (Figure S8). When calculating the percentage of induced CFUs that survived and became edited, the best results were seen for targets NS1 and NS4, followed by NS3 (Figure 4b). For NS2, no edited colonies were found (Figure S8d), removing this target from further consideration.

Based on these results it was decided to combine targets NS1 and NS4 in a double-edit target vector, and add the less efficient but still functional NS3 to a triple-edit target vector. To construct the multiplex target vectors, first the individual sgRNAs were combined by Golden Gate assembly to form a single sgRNA-array, which was then combined with the separate donor DNA pieces for simultaneous ligation into the pPMQAK1-CRISPR/Cas9 base vector (Figure 5a). Again, the [B], [B]+LVA, and [E*](104) base vectors were all tested.

**Figure 5.**
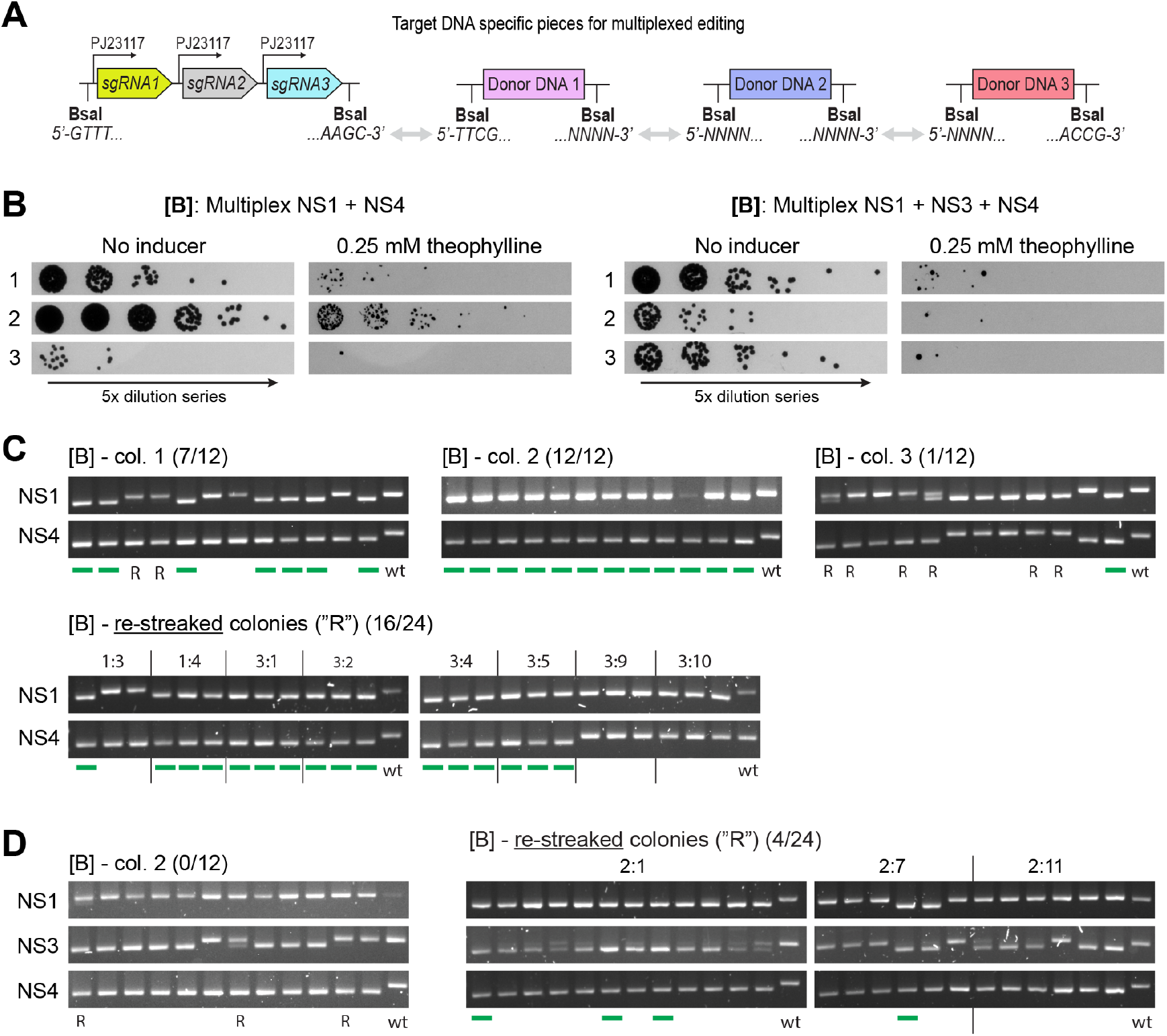
Inducible multiplexed CRISPR/Cas9 editing in S6803. (**A**) Schematic of the sgRNA and donor DNA pieces built for constructing a multi-target pPMQAK1-CRISPR/Cas9 target vector. The sgRNAs are supplied as a one-piece array, with the same overhangs as a single sgRNA. The donor DNA pieces are supplied as separate parts, with unique overhangs between the donors (*NNNN*), and the standard overhangs on the outermost ends. (**B**) Induction spot assay results for triplicate transformants of the [B] double- and triple-target constructs. 5x dilution series were plated on plates with or without 0.25 mM theophylline. (**C**) Editing results for the induced [B] double-target (NS1+NS4) construct. (**D**) Editing results for the induced [B] triple-target (NS1+NS3+NS4) construct. (**C-D**) A green line below a lane signals a fully segregated multi-edit in that screened colony. Colonies that appear segregated but have not been marked as such, are due to them having detectable wt-bands when the gels are more closely inspected. Fractions indicated the number of fully edited colonies out of the total number screened. A wt control shows how an unedited colony will appear. An “R” below a lane indicates a not fully segregated mutant that was re-streaked for a second round of induction.

Transformation of the multiplex target vectors into S6803 generally resulted in fewer transformants than seen previously for the single target vectors (Figure S9). The number of colonies was negatively correlated with the number of targets, and expression-strength of the CRISPR/Cas9 construct. Still, enough transformants were available that all constructs could be tested.

In a first test, pre-cultures of the transformants were prepared prior to induction. However, this pre-cultivation step appeared to select for cells that had mutated the CRISPR/Cas9 machinery, as most induced cells grew normally on theophylline and very few edits were found in the screened colonies (data not shown). A second attempt was done where the transformants were picked and plated directly on inducer plates (see method section). The representative spot assays now showed more promising results, with fewer cells surviving on the theophylline plates (Figure 5b & S10a). They also showed overall fewer surviving colonies for the triple-target (NS1+NS3+NS4) constructs, than the double-target (NS1+NS4) ones.

The induced transformants expressing the [B] (NS1+NS4) construct showed a large variation in editing efficiency (Figure 5c). For fully segregated colonies, the double-target editing efficiency ranged from 8-100%. Several screened colonies were edited only in one target or showed incomplete segregation; some of these latter ones could be pushed toward full segregation by re-streaking on new inducer-plates. It is likely that the segregation-resistant colonies had escaped the action of CRISPR/Cas9 by mutating it, thereby remaining un-edited but still viable. For the [B]+LVA (NS1+NS4) construct, re-streaking incompletely segregated colonies in a second round of induction ultimately yielded some double mutants (Figure S10b). However, for the [E*](104) (NS1+NS4) construct, while all screened colonies were fully segregated at the NS4-target (Figure S10b), there was no trace of editing in the NS1-target and a second round of induction was not attempted.

For the [B] (NS1+NS3+NS4) construct, only after a second round of induction were a few fully segregated triple-mutants identified (Figure 5d). Several more colonies were almost, but not yet fully segregated. The poor segregation of edits was observed mainly at the NS3-target site, which was known from the single-target data to be less amenable to editing than NS1 or NS4.

For the [B]+LVA (NS1+NS3+NS4) construct, edits were generally rare in any of the targets (Figure S10c); likely this weakest CRISPR/Cas9 construct was too weak to support this multi-editing attempt. For the [E*](104) (NS1+NS3+NS4) construct, the high survival on theophylline (Figure S10a) combined with few edits (Figure S10c) indicated that these had likely mutated the CRISPR/Cas9. Possibly the CRISPR/Cas9 activity for this [E*](104) constructs was too strong, causing cells to die from too rapidly occurring multiple DSBs or having already “escaped” the CRISPR/Cas9 by selecting against the effects of a leaky *cas9*-expression.

The different multiplexing success for the varyingly strong CRISPR/Cas9 constructs highlights the usefulness in having these options. Overall, the medium-strength construct [B] performed the best. Taken together, these results showed that multiplexed CRISPR/Cas9 editing using one single target vector is possible in S6803.

### Evaluating a self-targeting curing system to clear mutants of the CRISPR/Cas9 vector

An important feature of a CRISPR/Cas9 vector is that it should be easily cleared from cells after editing, leaving the mutant truly markerless. A common curing method is to grow edited cells without selecting for the vector, plate this, and screen colonies for plasmid loss. This method would likely be inefficient for the CRISPR/Cas9 vectors developed in this study, as RSF1010-replicon vectors are stably maintained in S6803 for prolonged periods under non-selective conditions.^46^ Another option is to add a counter-selection marker to the vector, such as *sacB* from *Bacillus subtilis* that causes sucrose-sensitivity.^47^ However, the *sacB*-method is only functional for glucose-tolerant wild type S6803 strains.^48^ For other S6803 wild types^49^ this curing method would be unusable.

Consideration this, a self-targeting sgRNA was added to the pPMQAK1-CRISPR/Cas9 vectors, enabling self-curing. To ensure that this curing sgRNA (named Cure-sgRNA) would not outcompete the main genome editing event and therefore mutate or clear the vector from the cells prematurely, it was purposefully designed to be weak. The alternative and weaker Cas9 NAG-PAM^13^ was used to find suitable protospacers on the pPMQAK1-backbone. The resulting Cure-sgRNAs were also constructed with truncated 3’-ends as this reduces DNA cleavage efficiency.^50^ In total, four Cure-sgRNAs, were individually evaluated in combination with the [B] and [E*](104) CRISPR/Cas9 vectors. The Cure-sgRNAs were expressed from the nickel- and cobalt-inducible *nrsD* promoter^51^ (Figure 6a); as using the constitutive P_BBa_J23117_ caused construct [E*](104) to loose its editing ability (data not shown). P_*nrsD*_ is native to S6803, so the coupled transcriptional regulator InrS is encoded from its own genome.^52^ P_*nrsD*_ has the lowest basal expression level of the characterized metal-inducible promoters from S6803.^53^ Its induced expression is estimated to be 2-fold higher than P_BBa_J23117_.^40,53,54^

**Figure 6.**
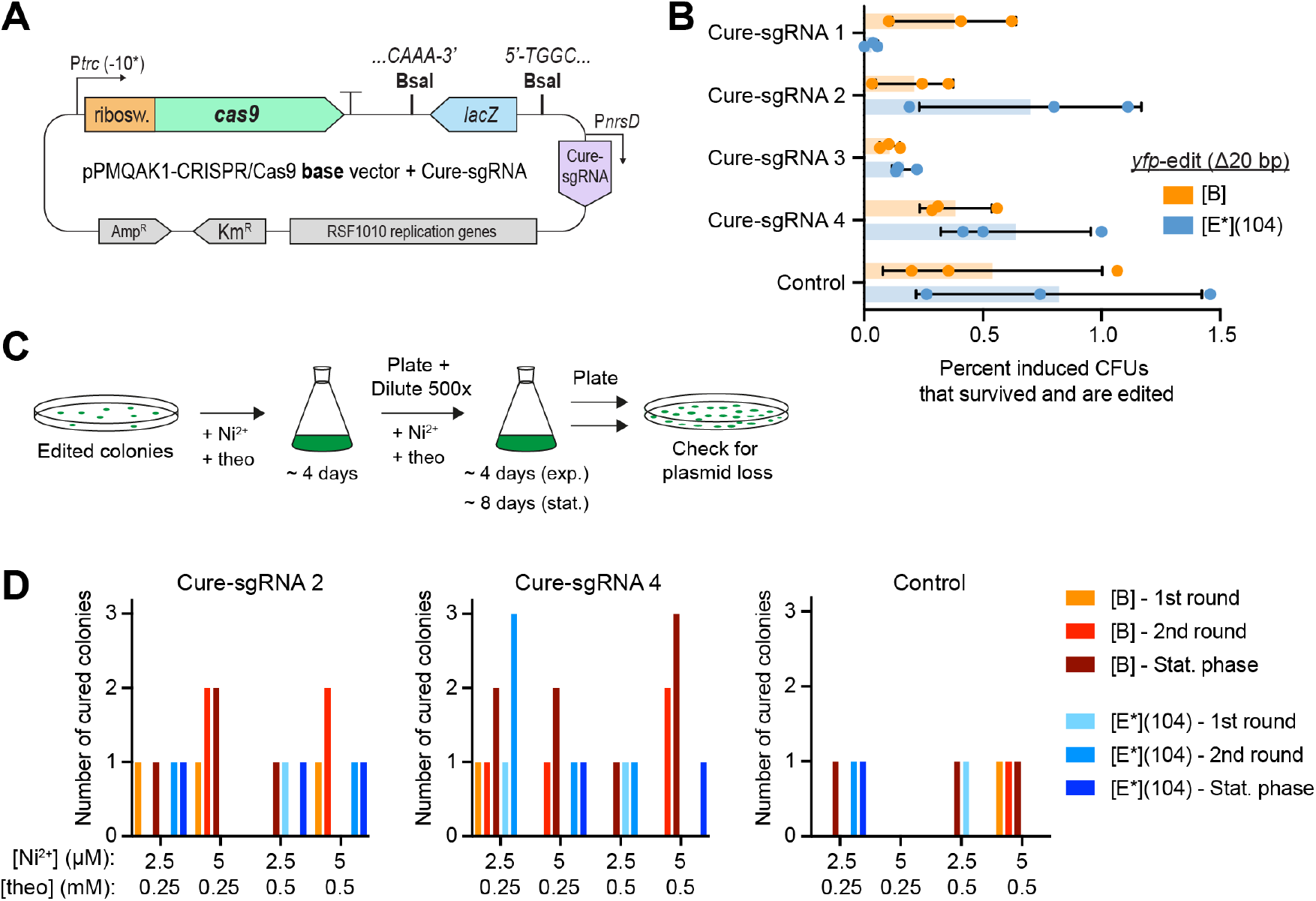
Evaluating an inducible curing system for the pPMQAK1-CRISPR/Cas9 vector. (**A**) Schematic showing the pPMQAK1-CRISPR/Cas9 base vector supplemented with a self-targeting Cure-sgRNA, expressed from the nickel- and cobalt-inducible P_*nrsD*_. (**B**) Percentage of induced CFUs that survived and became edited; for the *yfp*-targeting (Δ20 bp) constructs [B] and [E*](104) supplemented with P_*nrsD*_-Cure-sgRNA 1-4. The “Control” denotes a CRISPR/Cas9 vector without a Cure-sgRNA. Bars show the averages ± SD from biological triplicates, individual values are also shown. (**C**) Workflow for testing the curing system. Edited colonies were grown for two 4-day rounds in non-selective BG11 (-Km) with nickel and theophylline (“theo”). The second round was also grown until stationary phase. Plating of cells was done at the specified time-points, resulting in colonies that were screened for plasmid loss. (**D**) Results from testing the curing-ability of P_*nrsD*_-Cure-sgRNA 2 and 4 in NS1-edited (Δ30 bp) cells. The controls were edited cells harboring a CRISPR/Cas9 vector without a Cure-sgRNA. The curing efficiencies for the time-points specified in (C) are shown. Two different concentrations for nickel (2.5 vs. 5 µM) and theophylline (0.25 vs. 0.5 mM) were tested. Curing was determined both by screening colonies for the *kmR*-gene found on the pPMQAK1-backbone and growth on Km-plates. Each bar shows the total number of cured colonies out of 24 screened, i.e. 12 each for treated duplicate mutants.

The P_*nrsD*_-Cure-sgRNA supplemented CRISPR/Cas9 vectors were tested on the *yfp* (Δ20 bp) target as described previously. This target was revisited as it resulted in only moderate editing efficiencies, 42% and 54% for [B] and [E*](104), respectively (Figure 2d). The aim was to test if adding the self-targeting Cure-sgRNAs would negatively affect the efficiency of this suboptimal target editing. As P_*nrsD*_ is induced by cobalt in addition to nickel,^52^ all cobalt was omitted from the BG11 used in these experiments. Transformation results were not negatively affected by this change (Figure S11a), but it did affect the number of surviving colonies after CRISPR/Cas9-induction on theophylline (Figure 11b). Fewer or more surviving colonies were seen on regular BG11 for constructs [B] and [E*](104), respectively. The reason for this discrepancy is not known. Despite this, edited *yfp* (Δ20 bp) colonies were found for both the [B] and [E*](104) constructs variants (Figure S11c). However, editing efficiencies were consistently lower for constructs with a Cure-sgRNA (Figure S11c), also the number of surviving colonies on full-size inducer-plates varied (data not shown). To better judge the overall impact of adding the Cure-sgRNAs to the CRISPR/Cas9 vectors, the total percentage of induced CFUs that survived and became edited was calculated for each construct as described previously. The results showed a clear distinction among the stronger [E*](104) constructs (Figure 6b); the ones with Cure-sgRNA nr 1 or nr 3 performed worse than the control (no Cure-sgRNA). For the weaker [B] constructs, Cure-sgRNA nr 3 performed the poorest. Overall, the presence of Cure-sgRNA nr 2 and nr 4 had the least negative impact on genome editing.

The P_*nrsD*_-Cure-sgRNA supplemented CRISPR/Cas9 vectors were also tested on the NS1 (Δ30 bp) target (Figure S12), which previously resulted in high editing efficiencies of 92% and 100% for [B] and [E*](104), respectively (Figure S8b). These high editing efficiencies were maintained for the Cure-sgRNA constructs (Figure S12c). However, Cas9-induction on non-selective, regular BG11-plates (Figure S12b) showed that this high editing efficiency could not be combined with direct curing (Figure S13).

Finally, it was tested if induction of P_*nrsD*_-Cure-sgRNA could cure the pPMQAK1-CRISPR/Cas9 vector from edited cells. Based on previous data (Figure 6b), only constructs with Cure-sgRNA nr 2 and nr 4 were considered. Edited colonies were treated according to the workflow in Figure 6c. To control for spontaneous plasmid loss, edited colonies with vectors lacking a Cure-sgRNA were included. The curing extent was examined after each of two cultivation rounds; where cells were still exponentially growing (OD_730_ 0.2-1.2), and after the last round had reached stationary phase (OD_730_ > 10). Two concentrations were tested for the nickel to induce P_*nrsD*_ (2.5 vs. 5 µM) and theophylline to induce the CRISPR/Cas9 (0.25 vs. 0.5 mM). Overall, the curing results were better for the Cure-sgRNA constructs, compared to the controls (Figure 6d). There was no clear difference between the tested inducer concentrations, but a slight positive correlation with the longer cultivation time. Taking this into account, the total curing efficiencies reached after 4 days into the second cultivation round for construct [B] was 4.2%, 4.2%, and 1% for Cure-sgRNA nr 2, nr 4, and control, respectively. For construct [E*](104) it was 2.1%, 5.2%, and 1% for Cure-sgRNA nr 2, nr 4, and control, respectively. The Cure-sgRNA nr 4 worked the best, but only with a small margin. For these vector variants one would have to screen roughly 25 colonies to find a cured one, which is a 4-fold improvement to the 1% curing for the control constructs. To reduce the chemical stress imposed on the cells, the lower concentrations of nickel and theophylline are recommended.

## Conclusion

In this study we designed a riboswitch-based, theophylline-inducible CRISPR/Cas9-system for easy use in S6803. This system forgoes the common issue of Cas9-toxicity, enabling one to attain high transformation efficiencies for the pPMQAK1-CRISPR/Cas9 target vector. Inducing CRISPR/Cas9 in obtained transformants allowed edited colonies to be isolated. Single edits were fully segregated after just one passage on the inducer, reducing the editing time. The system was also shown to support multiplexed editing in S6803, enabling simultaneous editing of up to three targets using one single CRISPR/Cas9 vector. The [B] construct was most successful overall, supporting all attempted single and multiplexed edits. Although, the stronger [E*](104) construct could be useful in the case of weak sgRNAs that cannot be re-designed due to e.g. specific target site requirements. The two-step workflow, where transformation and CRISPR/Cas9 genome editing are done separately, was most reliable throughout this study. However, in some cases (e.g. adding the FLAG-tag to *rbcL*), edited colonies could be found among transformants that survived induction directly after transformation. Finally, a nickel-inducible curing system was added to, and evaluated for the [B] and [E*](104) CRISPR/Cas9 vector variants. The self-targeting Cure-sgRNA nr 4 was deemed best by enabling a 4-fold improvement in curing of edited cells.

## Methods

### Strains and general growth conditions

See Table S1 for strains used in this study. The S6803 wild type (a gift from Martin Fulda, University of Goettingen) is a non-motile GT-S derivative. A Δ*slr1181*::P_*psbA2*_-Yfp-B0015-Sp^r^ strain was used for *yfp* genome editing. All cultivations were done at 30 °C, 1% (v/v) CO_2_, and 30 (plates) or 50 (liquid) µE s^−1^ m^2^, using a Percival Climatics SE-1100 climate chamber. The BG11 media was buffered to pH 7.9 with 25 mM HEPES. For growth on solid media, 1.5% (w/v) agar and 0.3% (w/v) sodium thiosulphate was added to the BG11. When needed, antibiotics were supplemented (40 µg/ml kanamycin, 40 µg/ml spectinomycin). Growth in liquid media was monitored by OD_730_. Special conditions for e.g. CRISPR/Cas9-induction or plasmid curing are explained in the specific method sections below.

### Vector construction

For vectors used and constructed in this study see Table S1. All primers are listed in Table S2. All sub-cloning was done in *Escherichia coli* XL1-Blue.

The pPMQAK1-T vector^40^ has BpiI sites ready for Golden Gate cloning.^55^ The Gfp-reporter, and CRISPR/Cas9 base vectors were built by amplifying the required inserts, which added overhangs with compatible BpiI sites, and performing multi-piece assembly into pPMQAK1-T (see more details below). All Golden Gate cloning was done using Thermo Fisher Scientific T4 DNA ligase and FastDigest BpiI or BsaI (as specified). All Golden Gate cloning primers were designed using Benchling.^56^

The sequences of the evaluated promoters, P_*conII*_ and P_*trc*_, and riboswitches (B, C, E*) can be seen in the supporting information. The combinations were ordered as “Ultramer”-oligos from IDT.

To build the Gfp-reporter plasmids, the *gfp* and terminator (BBa_B0015) was amplified as a single piece from a plasmid available in-lab. The promoter-riboswitch pieces were amplified from the “Ultramer”-templates.

Before constructing the CRISPR/Cas9 base vectors, the pPMQAK1-T vector and *cas9* gene were domesticated for use with BsaI and BpiI, respectively. This was done either by PCR-based site-directed mutagenesis^57^ or sub-cloning using Golden Gate assembly. To build the CRISPR/Cas9 base vectors the following pieces were amplified and inserted into pPMQAK1-T: the P_*trc*_-riboswitch pieces using the “Ultramer”-templates, the BpiI-free *cas9* gene, a BBa_B0015 terminator, and the *lacZ*-gene as found in the pPMQAK1-T with extra added BsaI sites on either side for future cloning use. For the promoterless-*cas9* control vector, the P_*trc*_-riboswitch piece was left out. The constructed pPMQAK1-CRISPR/Cas9 base vectors, with P_*trc*_-riboswitch (B, C, or E*), were used in site-directed mutagenesis^57^ to make variants with an LVA-tag on Cas9, and variants with a weaker -10-box in P_*trc*_.

For the P_*nrsD*_-Cure-sgRNA supplemented [B] and [E*](104) CRISPR/Cas9 base vectors; the P_*nrsD*_-Cure-sgRNA was added as an extra piece in the base vector assembly described above. P_*nrsD*_ was amplified from the S6803 genome and the truncated (+54) sgRNA scaffold^50^ was amplified from an sgRNA template; these were joint together by overlap-PCR using the spacer sequences that had been added as overhangs. See supporting information for a P_*nrsD*_-Cure-sgRNA example sequence.

The pPMQAK1-CRISPR/Cas9 target vectors were built by Golden Gate cloning; the sgRNA and donor DNA were simultaneously inserted using the BsaI sites in the selected CRISPR/Cas9 base vector. The BsaI-containing overhangs added to the 5’- and 3’-ends of the amplified sgRNA and donor DNA were kept constant, see supporting information for details.

The single target sgRNAs were constructed by overlap-PCR, using the compatible spacer sequences added as overhangs to the BBa_J23117 promoter piece and the Cas9-handle-*S. pyogenes*-terminator piece. An example sgRNA sequence can be seen in the supporting information. For the multiplex target vectors, the sgRNAs were first combined into an array by Golden Gate sub-cloning in a pMD19-backbone. This sgRNA-array construction was done as described by Li et al.^45^ The final pMD19-sgRNA-array plasmid contained the necessary BsaI sites flanking the array, so it was directly added to the target vector Golden Gate assembly. The sgRNA spacers in this study (Table S3) were designed using Benchling.^56^ Off-target binding analysis (Table S4) was done using Benchling^56^ and the CasOT software.^58^

The donor DNAs were prepared by overlap-PCR of the amplified homology arms. An exception was when inserting the P_*psbA2*_-Yfp-B0015 cassette into *slr0168*; here the two homology arms and cassette were supplied as three separate pieces. For the multiplex target vectors, the donor DNA for each site was prepared separately, with overhangs as shown in Figure 5a.

### Strain construction

All constructed pPMQAK1 vectors were transformed into S6803 by electroporation, see supporting information for details. Each transformation used 5-10 ml exponentially growing S6803 (OD_730_ 0.6-1), and 100-350 ng of vector DNA. Cells were recovered for 16-24 h, before plating on selective media. For CRISPR/Cas9 vectors the recovered cells were pelleted, resuspended in 460 µl BG11, whereby 200 µl each was plated on selective BG11-plates without or with 0.25 mM theophylline. For preparation and use of the 200 mM theophylline stock, see supporting information. In some cases (e.g. the *yfp* edit, and testing the P_*nrsD*_-Cure-sgRNA vectors), the remaining cell suspensions were used to make spot assays (10 µl spots). Note, cells transformed with P_*nrsD*_-Cure-sgRNA supplemented vectors were resuspended in and plated on cobalt-free BG11.

### Inducing CRISPR/Cas9 genome editing

Induction was done on solid media. BG11-plates were supplemented with 0.25 mM (or as indicated) theophylline. Cobalt-free BG11 was used for transformants with P_*nrsD*_-Cure-sgRNA supplemented vectors. Induction was done in one of two ways; transformants were picked into pre-cultures (∼ 2-3 ml, done in 24-deep well plates) and cultivated for 3-4 days before plated on inducer-plates, or transformant colonies were suspended in a small volume (∼80 µl) of BG11 (or cobalt-free BG11 for Cure-sgRNA constructs) and directly plated on inducer-plates. The latter method is recommended. The described pre-cultures or colony-suspensions were diluted before plating on inducer-plates. The prepared “base”-dilution was OD_730_ 0.2, from which a 5x dilution series was prepared. The dilution series were used to make spot assays (4 µl spots) on indicated plate types, and the remaining volume (40-60 µl) of selected dilutions (commonly 5x, 25x, 125x) was then plated on full-size inducer-plates, see supporting information for details. After 10-14 days, healthy-looking colonies were screened. If needed, colonies with un-segregated edits were re-streaked on new inducer-plates.

### Evaluating genome editing and screening mutants

After CRISPR/Cas9-induction, surviving healthy-looking colonies were screened by colony-PCR. The editing efficiency was the percentage of edited colonies found amongst the ones screened. The second quantification used, the percentage of induced CFUs that survived and also became edited, was calculated by relating the found editing efficiency to the percentage of induced CFUs that survived. This latter percentage was estimated from the spot assays prepared for every editing experiment; the total plated CFUs and surviving, healthy CFUs were estimated from the plates without or with theophylline, respectively.

For the RbcL-FLAG immunoblot, 15 µg protein (soluble fraction) was separated by SDS-PAGE, transferred onto a 0.45 µm PVDF-membrane, and analysed using the WesternBreeze Chromogenic anti-mouse Kit with a primary anti-FLAG (F4042, Sigma-Aldrich) antibody at a 4000x dilution. Cells were lysed by bead beating (200 µl beads, 10 min total, 1 min on/off, 4°) in lysis buffer (50 Tris-HCl, 150 mM NaCl, Roche complete EDTA-free protease inhibitor); centrifugation for 30 min at 4° and 21,000xg allowed isolation of the soluble fraction supernatant.

### Curing of pPMQAK1-CRISPR/Cas9 vectors

Edited, fully segregated colonies were picked into non-selective regular BG11 cultures (start OD_730_ 0.05), supplemented with nickel (2.5 µM vs. 5 µM, using a 50 mM NiSO_4_ stock) and theophylline (0.25 mM vs. 0.5 mM) in all possible combinations. The cultures (2 ml) were grown in 24-deep well plates, sealed with gas permeable seals. Cultures were grown for 4 days, diluted 500x into new cultures with the same inducer concentrations, which were grown for a total of 8 days. Plating of cells on non-selective regular BG11-plates was done after both 4-day marks and the latter 8-day mark. Resulting colonies were screened for pPMQAK1-loss by checking for a retained kanamycin-resistance by colony-PCR and growth on Km-plates.

## Supporting information

Additional figures, tables, sequences of interest, and a general step-by-step method description for using the CRISPR/Cas9-system.

## Abbreviations

CRISPR: clustered regularly interspaced short palindromic repeats
Cas: CRISPR-associated
sgRNA: single chimeric guide RNA
PAM: protospacer adjacent motif
DSB: double-stranded break
HDR: homology-directed repair
RBS: ribosome binding site
CFU: colony-forming unit
wt: wild type
NS: neutral site

## Author Contribution

I. Cañadas, N.M. and E.P.H. conceived of the study, I. Cañadas contributed to the design of the study in an initial stage, I. Cengic and E.P.H. designed the rest and majority of the study, I. Cengic performed the experiments and analysed the data, I. Cengic wrote the manuscript, E.P.H. and N.M contributed to revising the manuscript.

## Acknowledgement

This work was funded by the European Union’s Horizon 2020 research and innovation program under Grant Agreement No. 760994 (ENGICOIN project), the Swedish Research Council (2020-04329), and Novo Nordisk Fonden (NNF20OC0061469).

## For Table of Contents Only

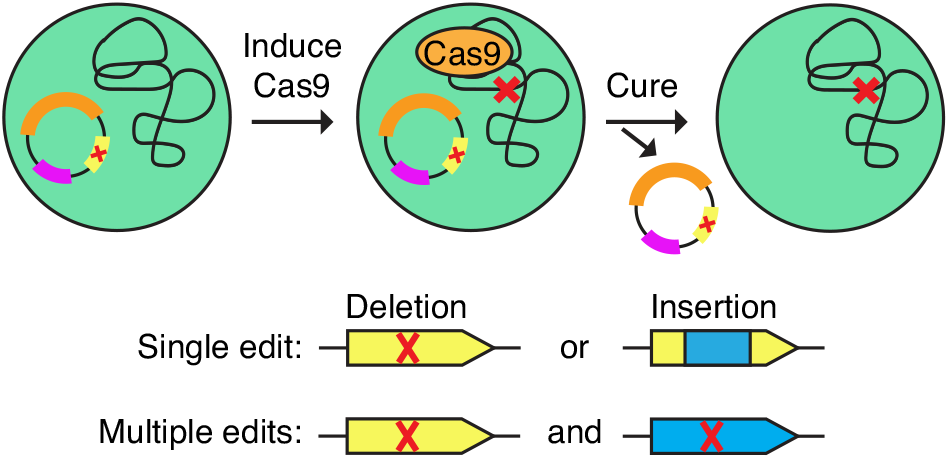

