## Supporting Information for "Inducible CRISPR/Cas9 allows for multiplexed and rapidly segregated single target genome editing in *Synechocystis* sp. PCC 6803"

**Contents:**

- Figure S1-S13
- Table S1-S4
- Sequences of promoters, riboswitches, and P*_nrsD_*-Cure-sgRNA
- Extra methods - Step-by-step protocol of using the CRISPR/Cas9-system

**Figures**

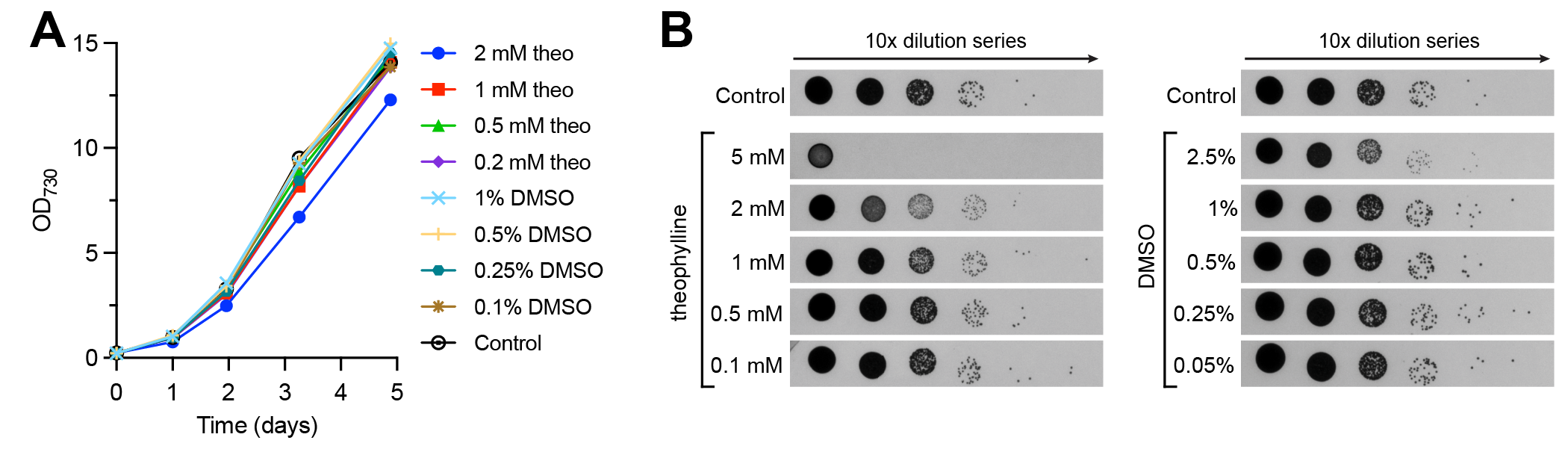

**Figure S1.** Evaluating the toxicity of theophylline towards S6803. (**A**) Single cultures were grown in BG11 supplemented with various concentrations of theophylline, or the respective amount of DMSO-carrier only. A control with neither was included. (**B**) Spot assay on BG11-plates supplemented with the indicated concentrations of theophylline or DMSO-carrier only. A control with neither was included. A wt S6803 culture was diluted to OD_730_ 0.2 and used to prepare a 10x dilution series that was plated (4 µl spots). Shown is representative data from two separate experiments.

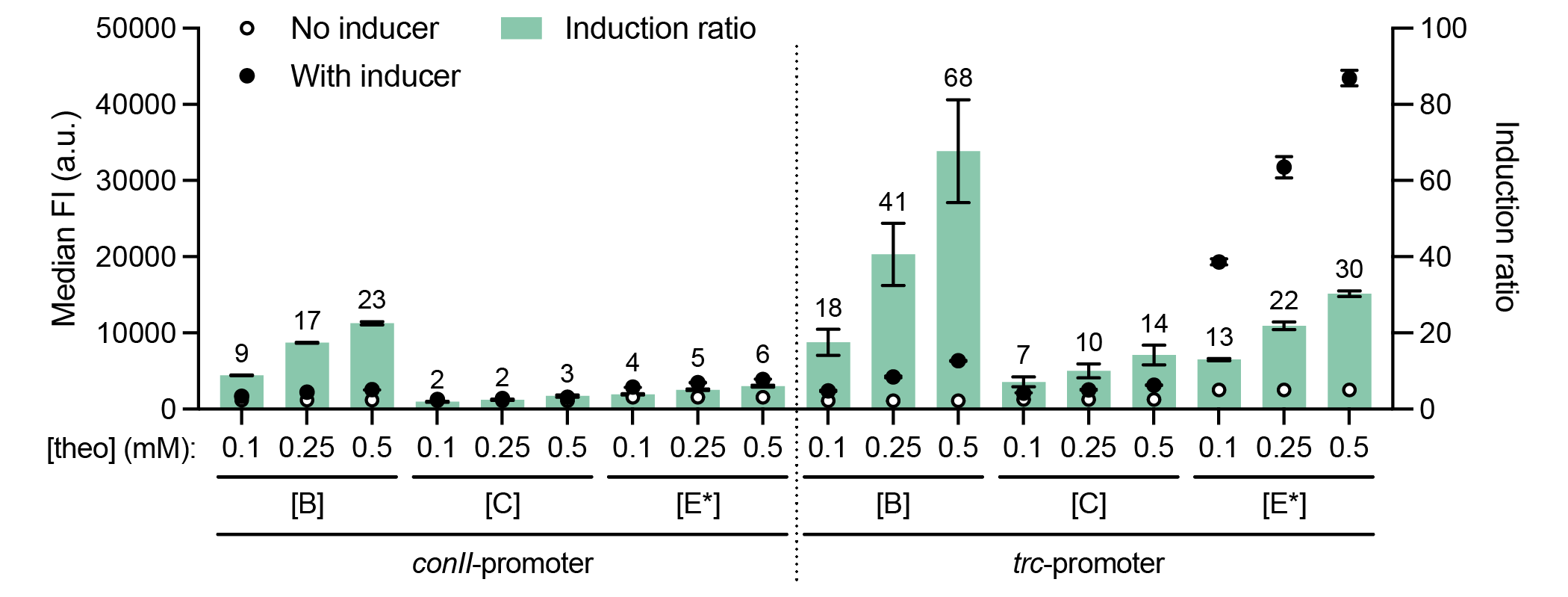

**Figure S2.** P*_conII_* and P*_trc_* in combination with riboswitches B, C, and E* were compared using a Gfp-reporter. Three non-toxic theophylline concentrations were tested: 0.1, 0.25, and 0.5 mM. Samples with no added inducer were supplemented with the respective amount of DMSO-carrier: 0.05%, 0.125%, and 0.25%. Fluorescence was measured 3 days after induction by flow cytometry (Beckman Coulter CytoFLEX, FITC-channel: emission 525 nm, excitation 488 nm). 10,000 events were acquired; data analysis was done using FlowJo (FlowJo LLC). The left y-axis shows the average median fluorescence intensity (filled or empty circle); the right y-axis shows the induction ratio (induced signal divided by un-induced signal, both values were first normalized against the wild type signal). The numbers above the bars are the calculated average induction ratios. All data is presented as averages ± SD from biological duplicates. Non-visible error-bars are smaller than the data symbol.

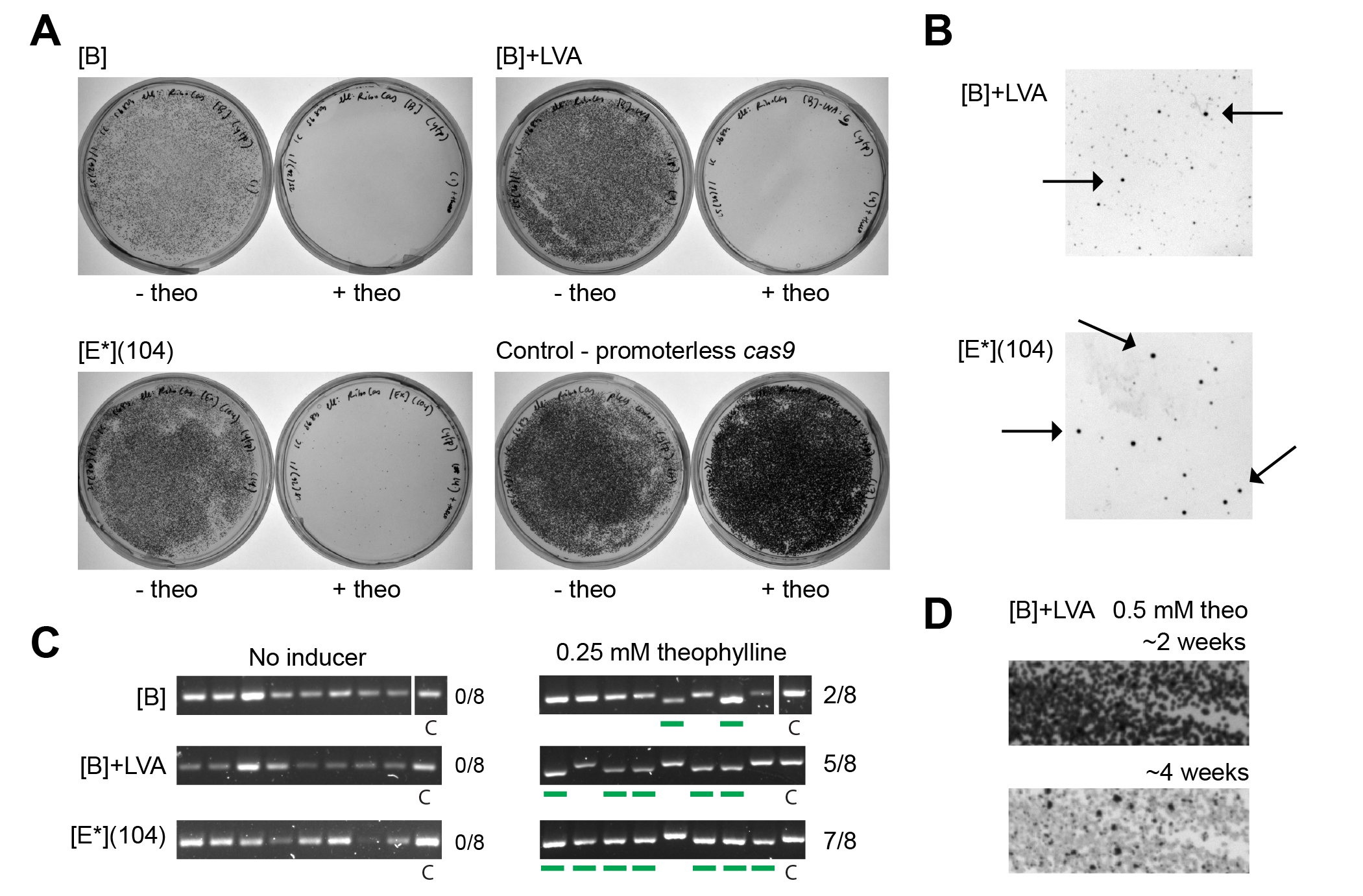

**Figure S3.** (**A**) Full plates of S6803 ∆*slr1181*::P*_psbA2_*-Yfp-B0015-Sp^r^, transformed with *yfp*-targeting (∆20 bp) pPMQAK1-CRISPR/Cas9 vector variants [B], [B]+LVA, [E*](104), and the control (no Cas9 expression). Plates without (-theo) and with 0.25 mM theophylline (+theo) are both shown. (**B**) Zoomed in sections of the +theo plates from (A) for construct variants [B]+LVA and [E*](104), showing surviving colonies. Arrows indicate examples of larger colonies, screened for editing. (**C**) Screening colonies from the transformation plates seen in (A), both from ones without and ones with inducer. A green line below a lane signals a fully edited (∆20 bp) mutant. A control (“C”) shows how an unedited colony will appear. Fractions indicated the number of fully edited colonies out of the total number screened. (**D**) Appearance of cells expressing the *yfp*-targeting (∆20 bp) [B]+LVA vector, after 2 and 4 weeks on 0.5 mM theophylline inducer.

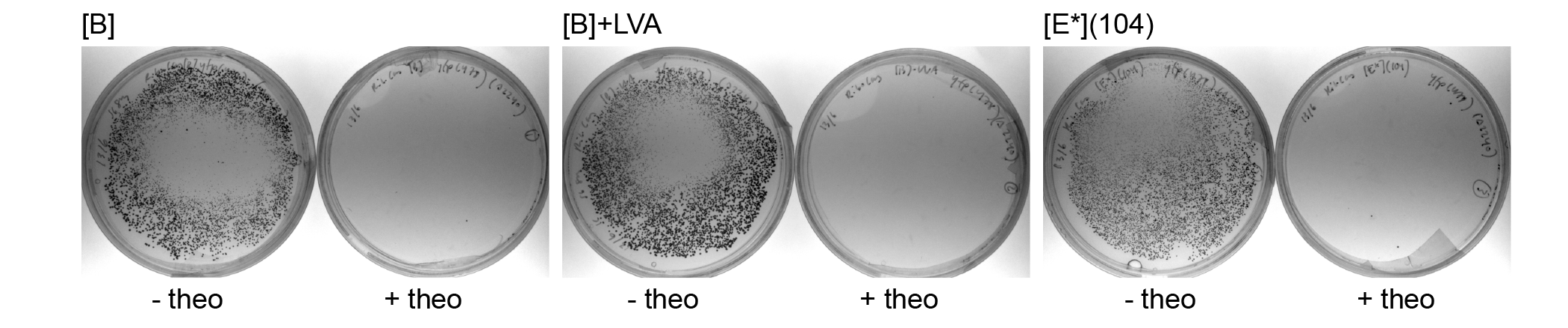

**Figure S4.** Full plates of S6803 ∆*slr1181*::P*_psbA2_*-Yfp-B0015-Sp^r^ transformed with *yfp*-targeting (∆2240 bp, whole Yfp-cassette) pPMQAK1-CRISPR/Cas9 vector variants [B], [B]+LVA, and [E*](104). Selective plates without (-theo) and with 0.25 mM theophylline (+theo) are both shown.

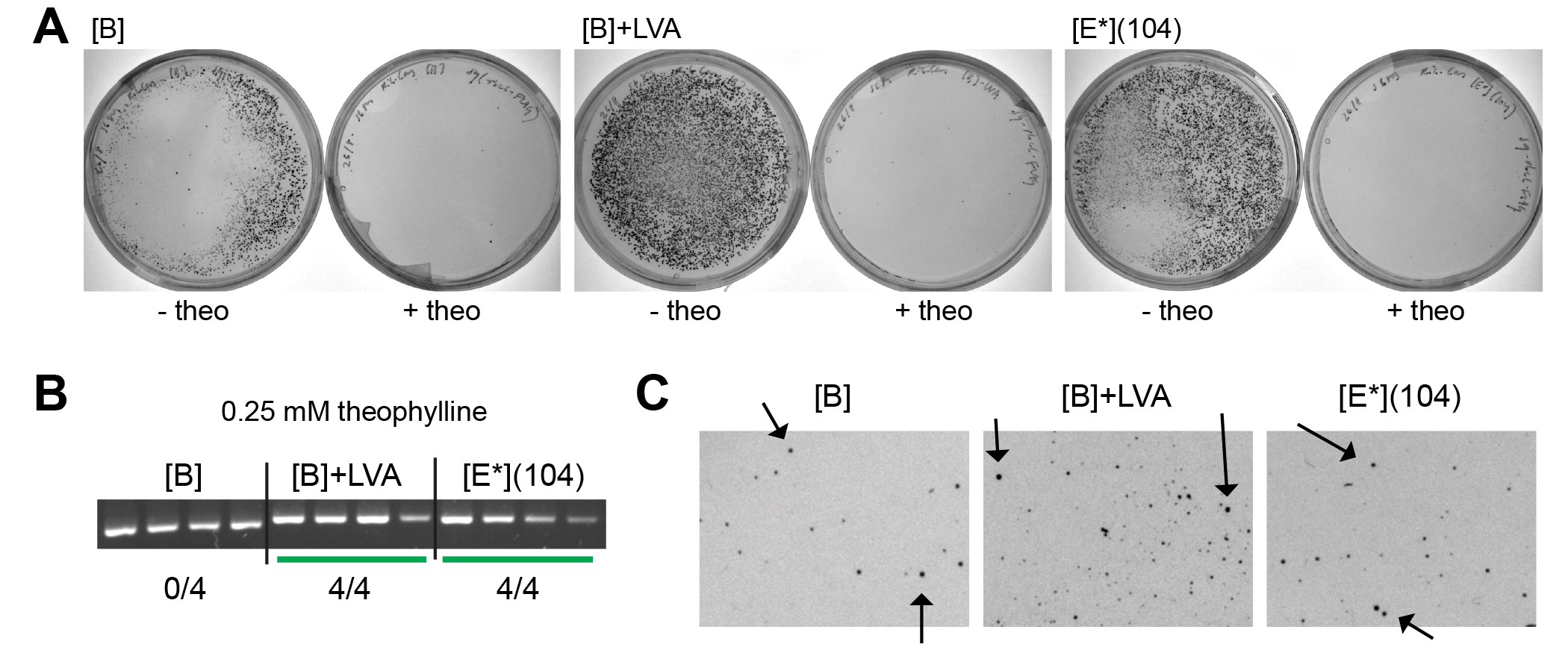

**Figure S5.** (**A**) Full plates of S6803 wt transformed with *rbcL*-targeting (adding a C-terminal FLAG) pPMQAK1-CRISPR/Cas9 vector variants [B], [B]+LVA, and [E*](104). Selective plates without (-theo) and with 0.25 mM theophylline (+theo) are both shown. (**B**) Screening surviving colonies from the +theo transformation plates seen in (A). A green line below a lane signals a fully edited (+24 bp) mutant. Fractions indicated the number of fully edited colonies out of the total number screened. (**C**) Zoomed-in sections of inducer-plates (0.25 mM), showing the appearance of different sized colonies. The larger ones (examples indicated by arrows) were screened for editing.

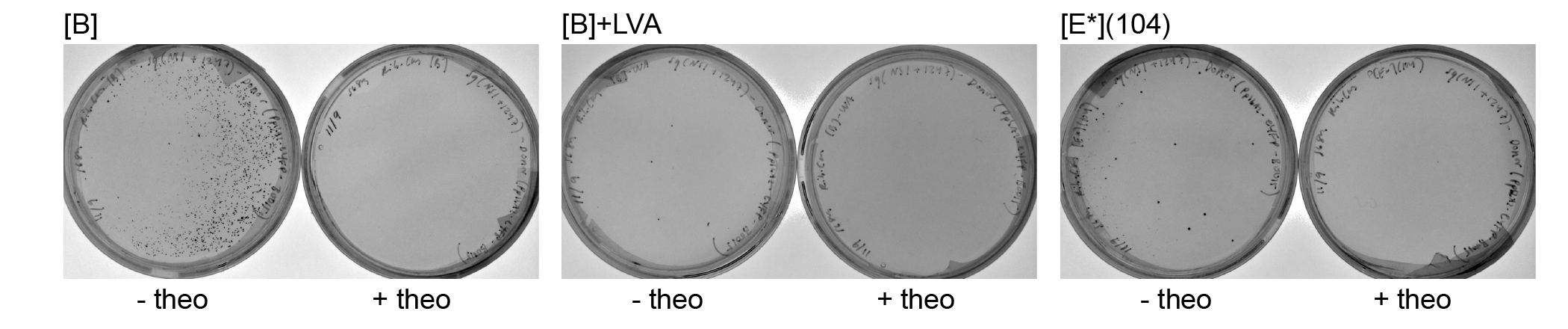

**Figure S6.** Full plates of S6803 wt transformed with *slr0168*-targeting (+1220 bp, P*_psbA2_*-Yfp-B0015 cassette) pPMQAK1-CRISPR/Cas9 vector variants [B], [B]+LVA, and [E*](104). Selective plates without (-theo) and with 0.25 mM theophylline (+theo) are both shown.

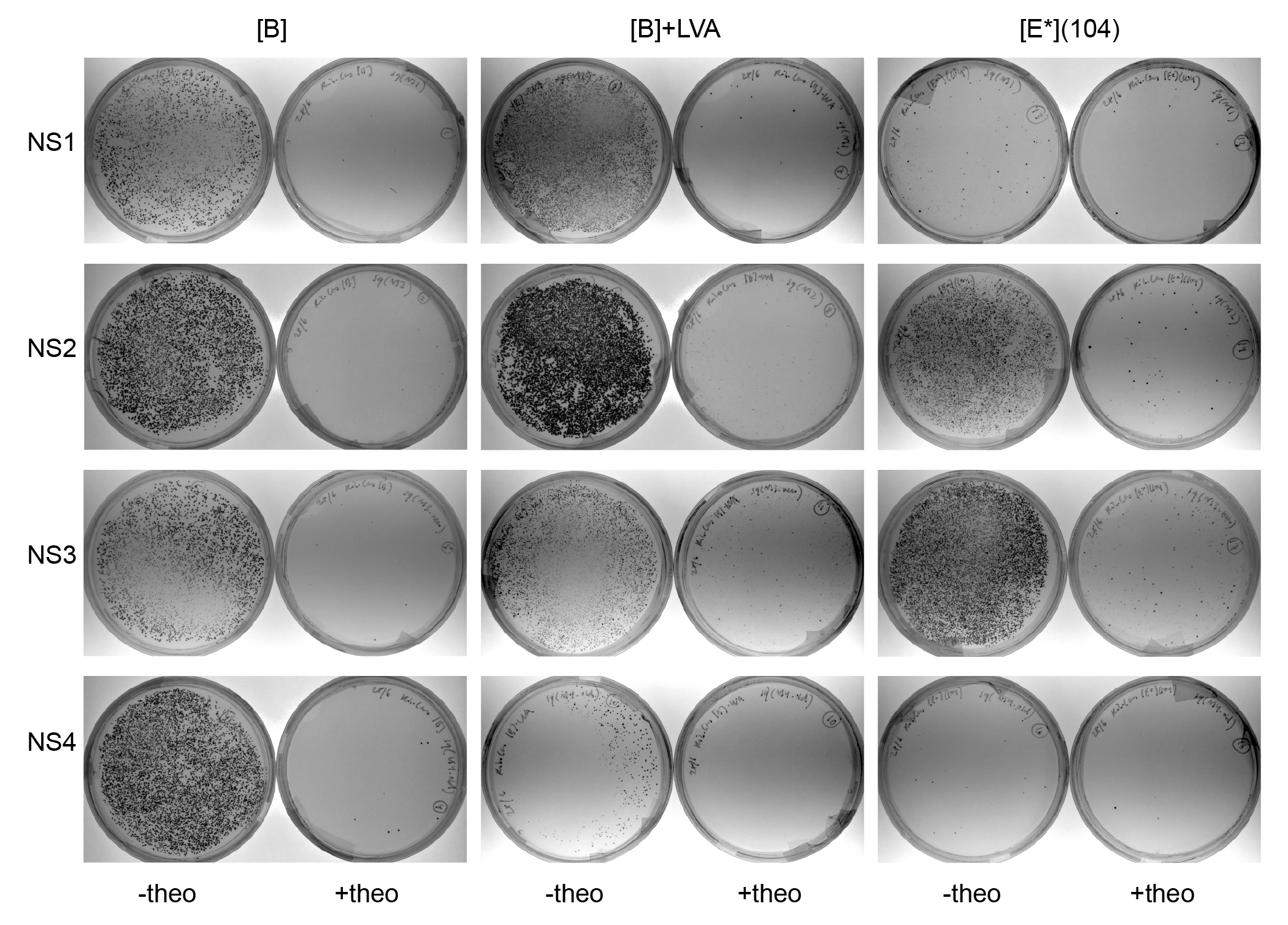

**Figure S7.** Full plates of S6803 wt transformed with the individually neutral site-targeting pPMQAK1-CRISPR/Cas9 vector variants [B], [B]+LVA, and [E*](104). Selective plates without (-theo) and with 0.25 mM theophylline (+theo) are both shown.

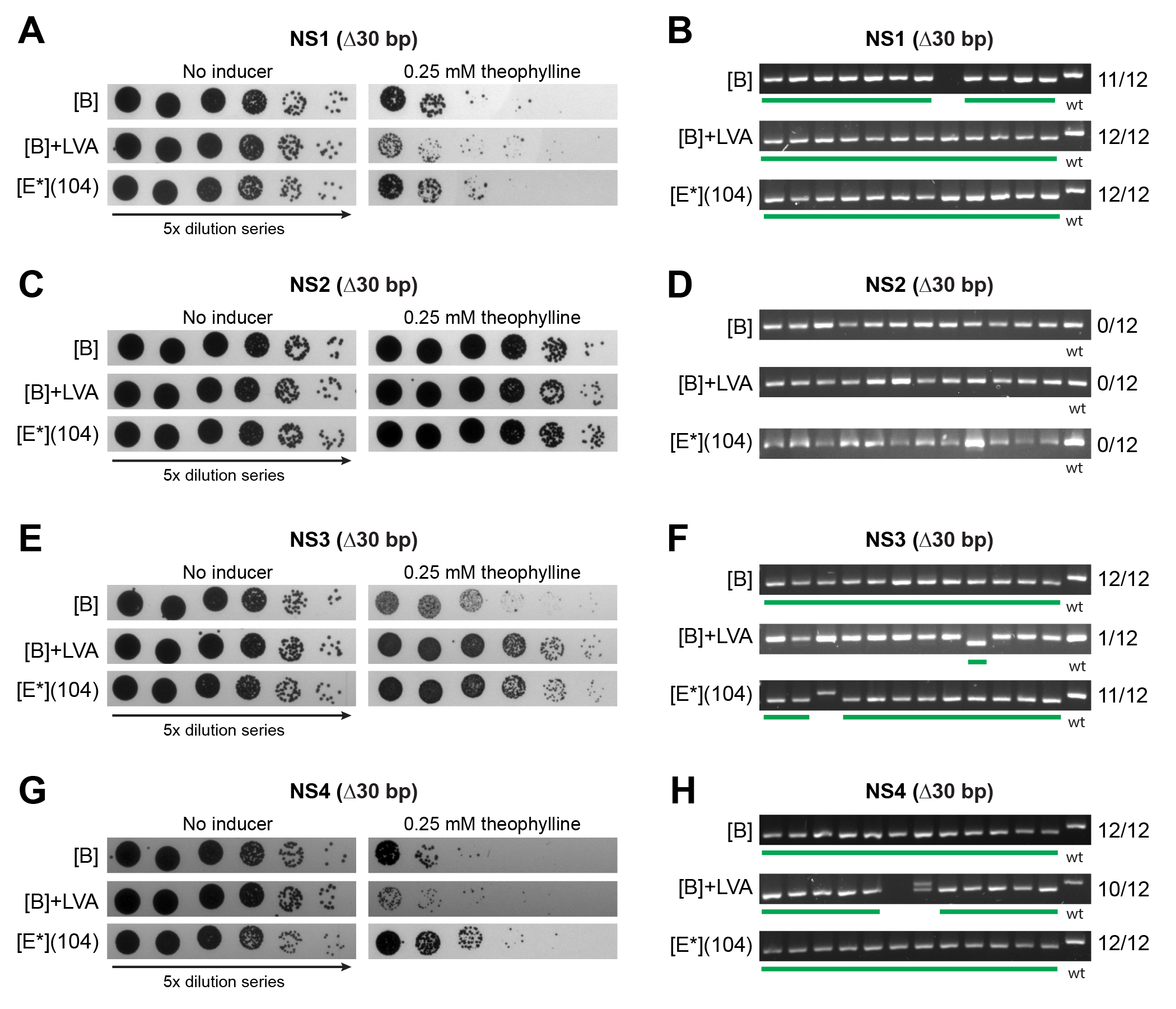

**Figure S8.** Results from induced editing (∆30 bp) of NS1-4 as single targets. (**A, C, E, G**) Induction spot assay results for the individual NS1-4 targets, with the indicated pPMQAK1-CRISPR/Cas9 vector variant. 5x dilution series were plated on plates without or with 0.25 mM theophylline. Done for biological duplicates, representative data is shown. (**B, D, F, H**) Editing results for the individual NS1-4 (∆30 bp) targets, with the indicated CRISPR/Cas9 vector variants. A green line below a lane signals a fully edited (∆30 bp) mutant. A wt control shows how an unedited colony will appear. Fractions indicated the number of fully edited colonies out of the total number screened.

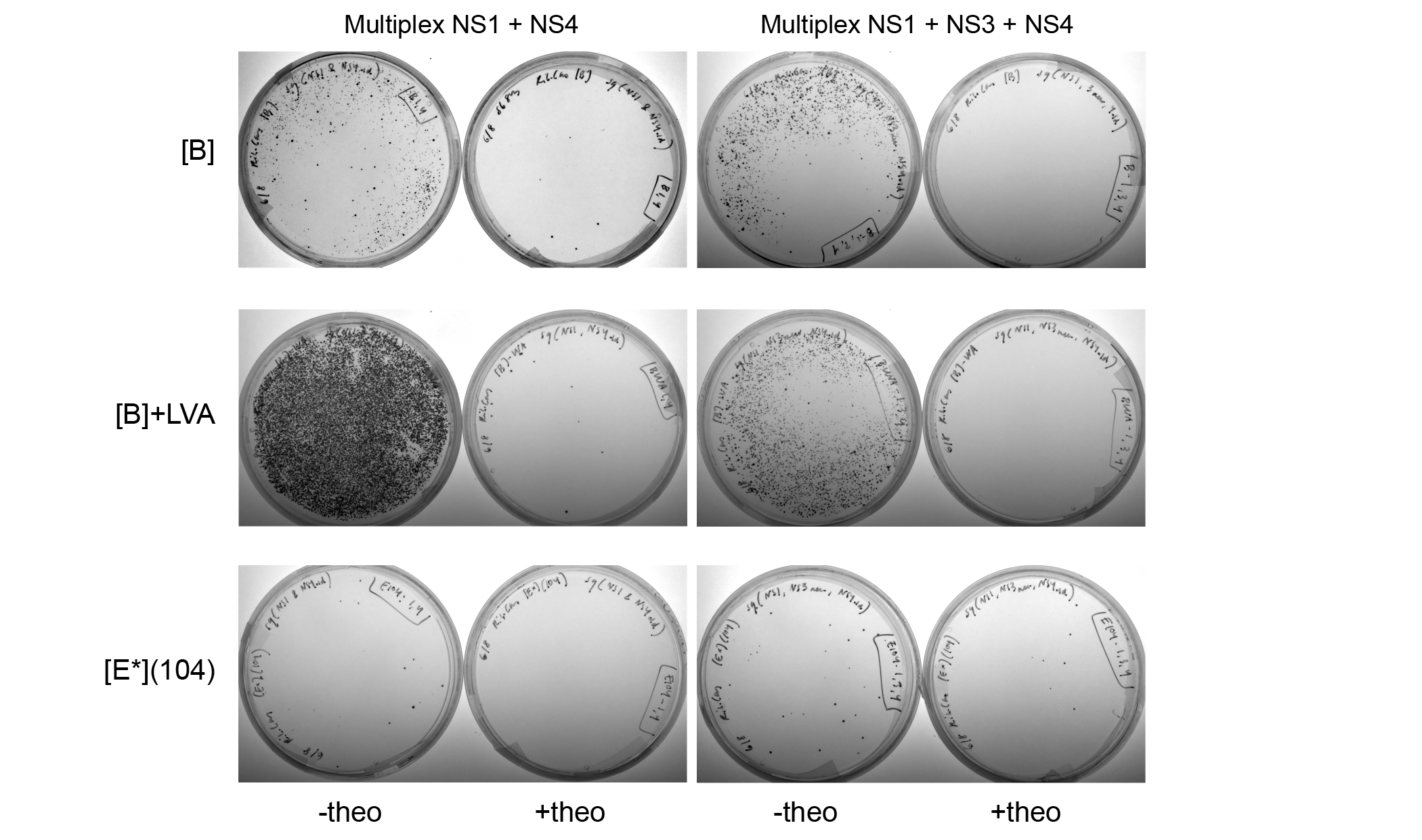

**Figure S9.** Full plates of S6803 wt transformed with multi-target pPMQAK1-CRISPR/Cas9 vector variants [B], [B]+LVA, and [E*](104). The multiplexed constructs targeted either NS1+NS4, or NS1+NS3+NS4. Selective plates without (-theo) and with 0.25 mM theophylline (+theo) are both shown.

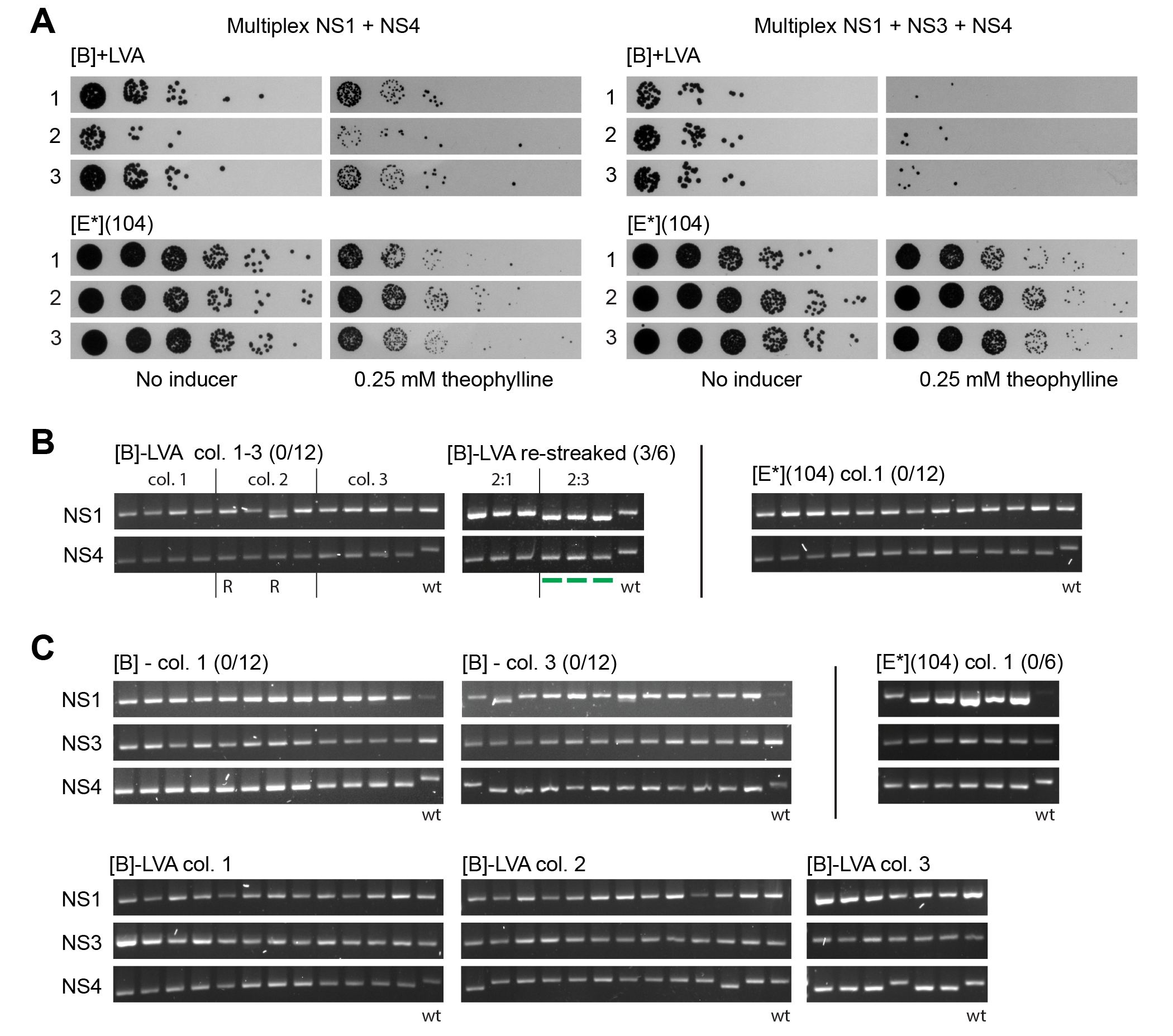

**Figure S10.** Additional results from induced multi-editing of the combined targets NS1+NS4 and NS1+NS3+NS4 in S6803. (**A**) Induction spot assay results for triplicate transformants of double- and triple-target pPMQAK1-CRISPR/Cas9 vector variants [B]+LVA and [E*](104). 5x dilution series were plated on plates with or without 0.25 mM theophylline. (**B-C**) Editing results for the (B) double-target (NS1+NS4) or (C) triple-target (NS1+NS3+NS4) constructs. The [B], [B]+LVA, and [E*](104) vector variants were screened where indicated. A green line below a lane signals a fully segregated multi-edit in that screened colony. Fractions indicated the number of fully edited colonies out of the total number screened. A wt control shows how an unedited colony will appear. An “R” below a lane indicates a not fully segregated mutant that was re-streaked for a second round of induction.

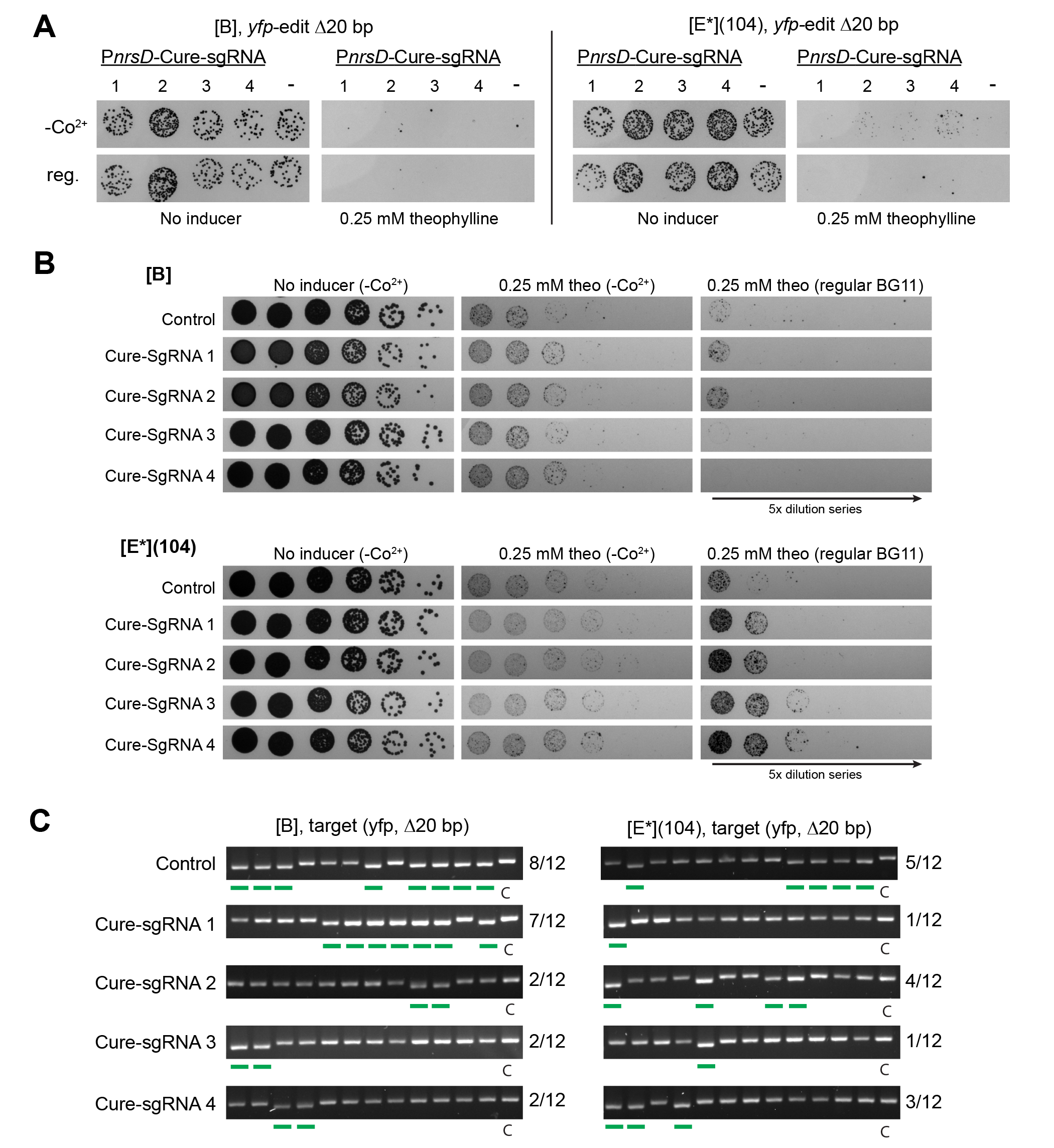

**Figure S11.** Data for constructs [B] and [E*](104) supplemented with P*_nrsD_*-Cure-sgRNA 1-4, targeting *yfp* (∆20 bp). (**A**) S6803 transformation results after plating on selective plates made with regular BG11 (“reg.”) or cobalt-free BG11, without or with 0.25 mM theophylline. The “-“ denotes vectors without a Cure-sgRNA. (**B**) Induction spot assay results. The “Control” denotes a CRISPR/Cas9 vector without a Cure-sgRNA. 5x dilution series were plated on cobalt-free BG11-plates without or with 0.25 mM theophylline, and also on inducer-plates made of regular BG11. Done for biological triplicates, representative data is shown. (**C**) Editing results for *yfp* (∆20 bp), for cells induced on cobalt-free BG11-plates. The “Control” denotes a vector without a Cure-sgRNA. A green line below a lane signals a fully edited (∆20 bp) mutant. A control (“C”) shows how an unedited colony will appear. Fractions indicated the number of fully edited colonies out of the total number screened.

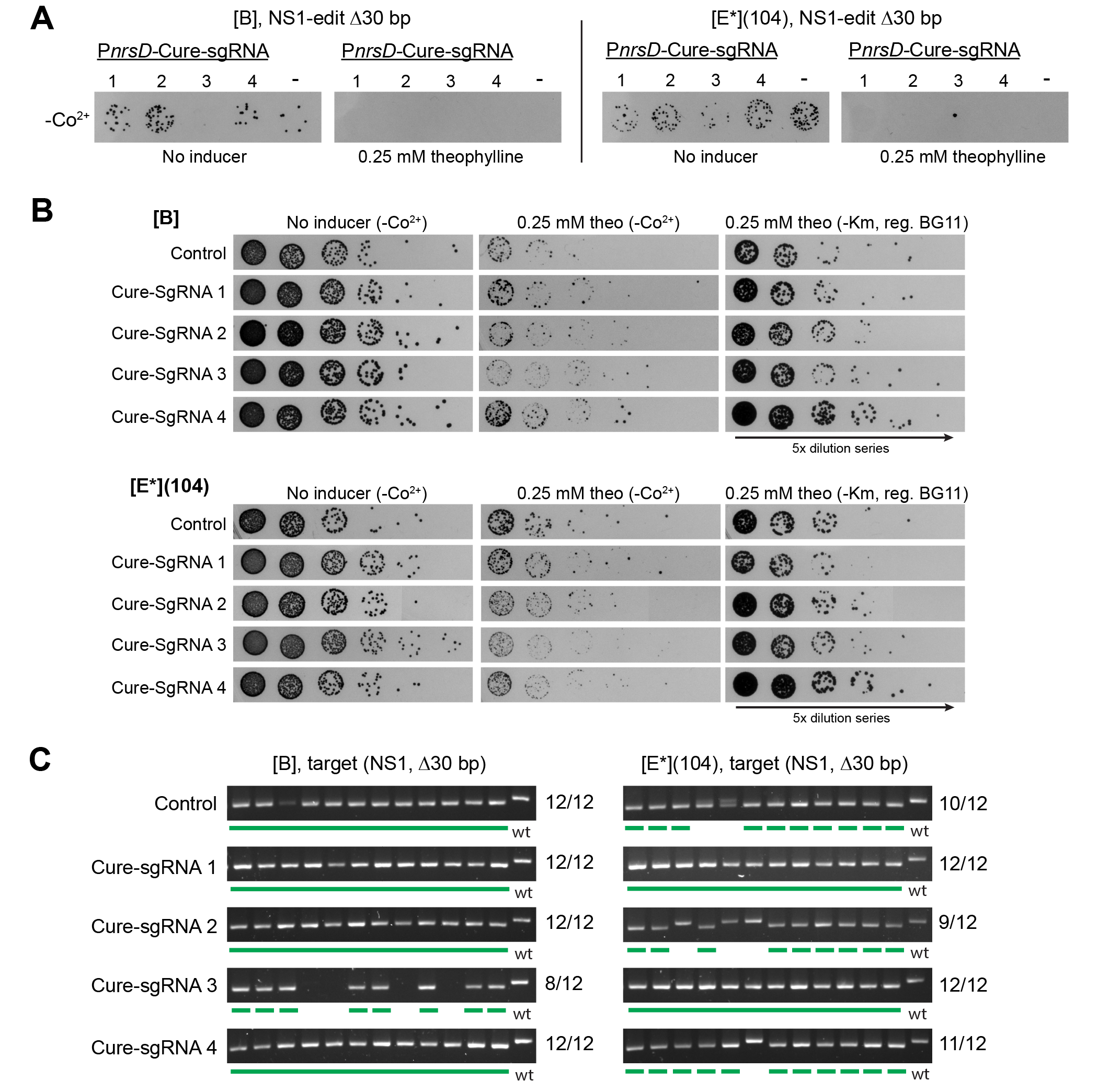

**Figure S12.** Data for constructs [B] and [E*](104) supplemented with P*_nrsD_*-Cure-sgRNA 1-4, targeting NS1 (∆30 bp). (**A**) S6803 transformation results after plating on selective plates made with cobalt-free BG11, without or with 0.25 mM theophylline. The “-“ denotes vectors without a Cure-sgRNA. (**B**) Induction spot assay results. The “Control” denotes a vector without a Cure-sgRNA. 5x dilution series were plated on cobalt-free BG11-plates without or with 0.25 mM theophylline inducer, and also on non-selective (-Km) regular BG11-plates with 0.25 mM theophylline. Done for biological triplicates, representative data is shown. (**C**) Editing results for NS1 (∆30 bp), for cells induced on cobalt-free, selective BG11-plates. The “Control” denotes a vector without a Cure-sgRNA. A green line below a lane signals a fully edited (∆30 bp) mutant. A wt control shows how an unedited colony will appear. Fractions indicated the number of fully edited colonies out of the total number screened.

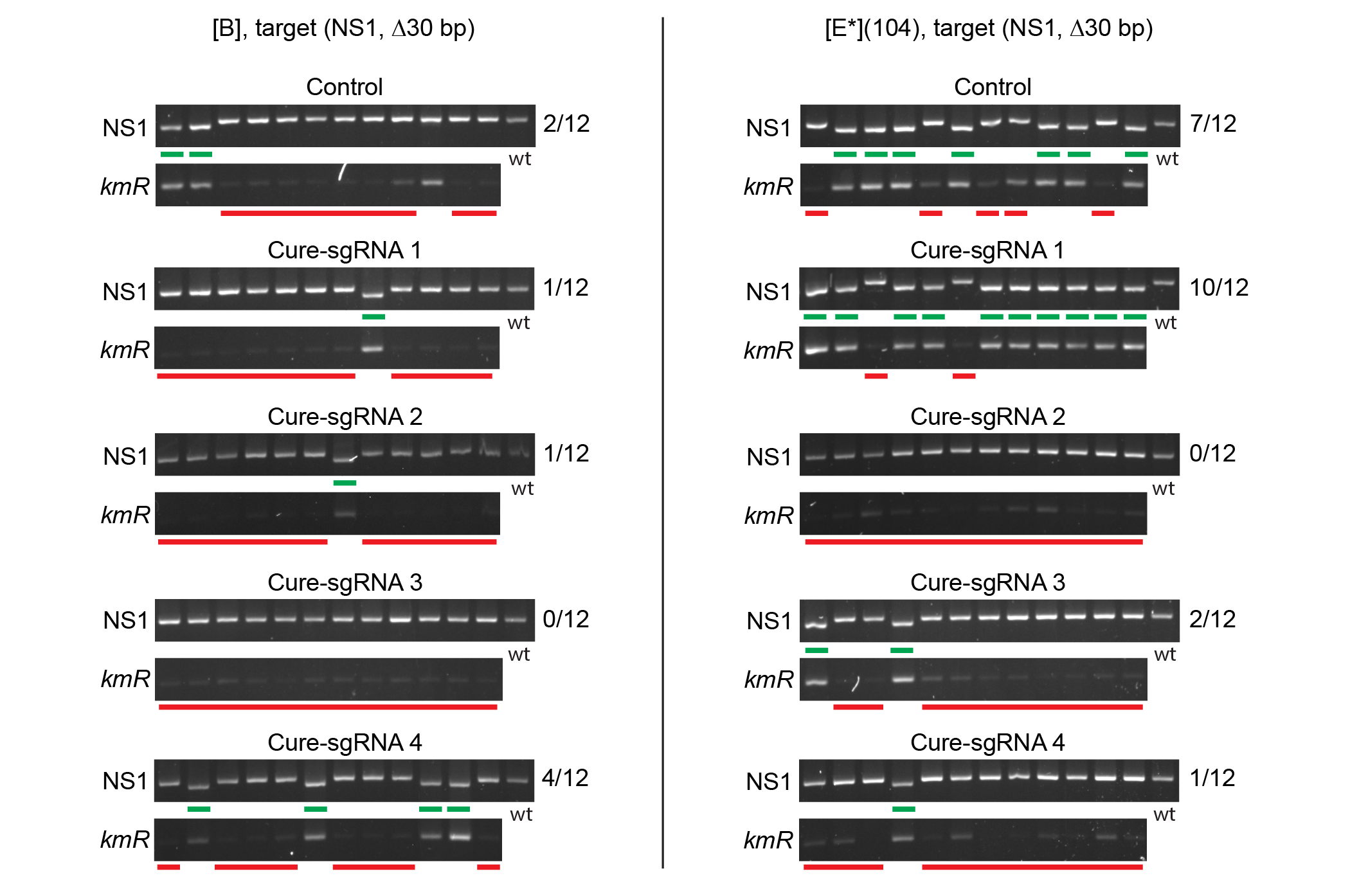

**Figure S13.** Editing results for construct [B] and [E*](104) supplemented with P*_nrsD_*-Cure-sgRNA 1-4, targeting NS1 (∆30 bp), induced on non-selective (-Km) regular BG11-plates with 0.25 mM theophylline. The “Control” denotes a vector without a Cure-sgRNA. Screening was done both for editing in NS1 (∆30 bp) and for presence of the pPMQAK1-CRISPR/Cas9 vector by screening for its kanamycin resistance gene (*kmR*). A green line below a lane signals an NS1-edited (∆30 bp) mutant. A wt control shows how an unedited colony will appear. Fractions indicated the number of fully edited colonies out of the total number screened. All screened colonies were also patched on selective Km-plates to check if they were still resistant. A red line below a lane signals that this colony did not manage to grow on the Km-plate.

**Tables**

**Table S1.** Strains and vectors used or constructed in this study. Only CRISPR/Cas9 **base** vectors are shown, the target vectors are not listed. (B0015 is terminator BBa_B0015).

| **Strain or Vectors** | **Relevant characteristics** | **Source** |
| --- | --- | --- |
| **Strains** |  |  |
| *Escherichia coli* XL1-Blue | Cloning host | Stratagene |
| *Synechocystis* sp. PCC 6803 |  |  |
| wild type | Non-motile, GT-S derivative | M. Fulda |
| ∆*slr1181*::P*_psbA2_*-Yfp-B0015-Sp^r^ | Yfp expressed from P*psbA2* in site *slr1181*, Sp^r^ | In-lab, D. Kaczmarzyk |
| **Vectors** |  |  |
| pPMQAK1-T | Replicative vector (RSF1010). Amp^r^, Km^r^ | Vasudevan et al.^1^ |
| pPMQAK1-T (no BsaI) | Replicative vector (RSF1010), BsaI site removed. Amp^r^, Km^r^ | This study |
| pPMQAK1-P*_conII_*-[B]-Gfp-B0015 | Gfp-reporter vector, Amp^r^, Km^r^ | This study |
| pPMQAK1-P*_conII_*-[C]-Gfp-B0015 | Gfp-reporter vector, Amp^r^, Km^r^ | This study |
| pPMQAK1-P*_conII_*-[E*]-Gfp-B0015 | Gfp-reporter vector, Amp^r^, Km^r^ | This study |
| pPMQAK1-P*_trc_*-[B]-Gfp-B0015 | Gfp-reporter vector, Amp^r^, Km^r^ | This study |
| pPMQAK1-P*_trc_*-[C]-Gfp-B0015 | Gfp-reporter vector, Amp^r^, Km^r^ | This study |
| pPMQAK1-P*_trc_*-[E*]-Gfp-B0015 | Gfp-reporter vector, Amp^r^, Km^r^ | This study |
| pPMQAK1-P*_trc_*-[B]-Cas9-B0015-*lacZ* | CRISPR/Cas9 base vector [B], Amp^r^, Km^r^ | This study |
| pPMQAK1-P*_trc_*-[C]-Cas9-B0015-*lacZ* | CRISPR/Cas9 base vector [C], Amp^r^, Km^r^ | This study |
| pPMQAK1-P*_trc_*-[E*]-Cas9-B0015-*lacZ* | CRISPR/Cas9 base vector [E*], Amp^r^, Km^r^ | This study |
| pPMQAK1-P*_trc_*-[B]-Cas9+LVA-B0015-*lacZ* | CRISPR/Cas9 base vector [B]+LVA, Amp^r^, Km^r^ | This study |
| pPMQAK1-P*_trc_*-[C]-Cas9+LVA-B0015-*lacZ* | CRISPR/Cas9 base vector [C]+LVA, Amp^r^, Km^r^ | This study |
| pPMQAK1-P*_trc_*-[E*]-Cas9+LVA-B0015-*lacZ* | CRISPR/Cas9 base vector [E*]+LVA, Amp^r^, Km^r^ | This study |
| pPMQAK1-P*_trc_*(-10:TATTGT)-[B]-Cas9-B0015-*lacZ* | CRISPR/Cas9 base vector [B](104), Amp^r^, Km^r^ | This study |
| pPMQAK1-P*_trc_*(-10:TATTGT)-[C]-Cas9-B0015-*lacZ* | CRISPR/Cas9 base vector [C](104), Amp^r^, Km^r^ | This study |
| pPMQAK1-P*_trc_*(-10:TATTGT)-[E*]-Cas9-B0015-*lacZ* | CRISPR/Cas9 base vector [E*](104), Amp^r^, Km^r^ | This study |
| pPMQAK1-P*_trc_*(-10:GACTAT)-[B]-Cas9-B0015-*lacZ* | CRISPR/Cas9 base vector [B](116), Amp^r^, Km^r^ | This study |
| pPMQAK1-P*_trc_*(-10:GACTAT)-[C]-Cas9-B0015-*lacZ* | CRISPR/Cas9 base vector [C](116), Amp^r^, Km^r^ | This study |
| pPMQAK1-P*_trc_*(-10:GACTAT)-[E*]-Cas9-B0015-*lacZ* | CRISPR/Cas9 base vector [E*](116), Amp^r^, Km^r^ | This study |
| pPMQAK1-P*_trc_*(-10:GATTGT)-[B]-Cas9-B0015-*lacZ* | CRISPR/Cas9 base vector [B](117), Amp^r^, Km^r^ | This study |
| pPMQAK1-P*_trc_*(-10:GATTGT)-[C]-Cas9-B0015-*lacZ* | CRISPR/Cas9 base vector [C](117), Amp^r^, Km^r^ | This study |
| pPMQAK1-P*_trc_*(-10:GATTGT)-[E*]-Cas9-B0015-*lacZ* | CRISPR/Cas9 base vector [E*](117), Amp^r^, Km^r^ | This study |
| pPMQAK1-Cas9-B0015-*lacZ* | Control vector, Cas9 lacks promoter and ATG, Amp^r^, Km^r^ | This study |
| pPMQAK1-P*_trc_*-[B]-Cas9-B0015-*lacZ*-P*_nrsD_*-Cure-sgRNA 1 | CRISPR/Cas9 base vector [B], inducible Cure-sgRNA 1, Amp^r^, Km^r^ | This study |
| pPMQAK1-P*_trc_*-[B]-Cas9-B0015-*lacZ*-P*_nrsD_*-Cure-sgRNA 2 | CRISPR/Cas9 base vector [B], inducible Cure-sgRNA 2, Amp^r^, Km^r^ | This study |
| pPMQAK1-P*_trc_*-[B]-Cas9-B0015-*lacZ*-P*_nrsD_*-Cure-sgRNA 3 | CRISPR/Cas9 base vector [B], inducible Cure-sgRNA 3, Amp^r^, Km^r^ | This study |
| pPMQAK1-P*_trc_*-[B]-Cas9-B0015-*lacZ*-P*_nrsD_*-Cure-sgRNA 4 | CRISPR/Cas9 base vector [B], inducible Cure-sgRNA 4, Amp^r^, Km^r^ | This study |
| pPMQAK1-P*_trc_*(-10:TATTGT)-[E*]-Cas9-B0015-*lacZ*-P*_nrsD_*-Cure-sgRNA 1 | CRISPR/Cas9 base vector [E*](104), inducible Cure-sgRNA1, Amp^r^, Km^r^ | This study |
| pPMQAK1-P*_trc_*(-10:TATTGT)-[E*]-Cas9-B0015-*lacZ*-P*_nrsD_*-Cure-sgRNA 2 | CRISPR/Cas9 base vector [E*](104), inducible Cure-sgRNA2, Amp^r^, Km^r^ | This study |
| pPMQAK1-P*_trc_*(-10:TATTGT)-[E*]-Cas9-B0015-*lacZ*-P*_nrsD_*-Cure-sgRNA 3 | CRISPR/Cas9 base vector [E*](104), inducible Cure-sgRNA3, Amp^r^, Km^r^ | This study |
| pPMQAK1-P*_trc_*(-10:TATTGT)-[E*]-Cas9-B0015-*lacZ*-P*_nrsD_*-Cure-sgRNA 4 | CRISPR/Cas9 base vector [E*](104), inducible Cure-sgRNA4, Amp^r^, Km^r^ | This study |
| pMD19-(EcoRI, XbaI)-Sp^r^ | Vector with cloning sites, for building sgRNA-array template vectors (below), Sp^r^ | In-lab, L. Yao |
| pMD19-(EcoRI, XbaI)-Cm^r^ | Vector with cloning sites, for building sgRNA-array template vectors (below), Cm^r^ | In-lab, L. Yao |
| pMD19-BsaI-P_BBa_J23117_-sgRNA-BsaI-Sp^r^ | Template vector – sgRNA single and multiplex-array (as Li et al.),^2^ flanking BsaI-sites added, Sp^r^ | This study |
| pMD19-Cas9_handle-S.pyogenes_terminator-P_BBa_J23117_-Cm^r^ | Template vector – sgRNA multiplex-array (as Li et al.),^2^ Cm^r^ | This study |

**Table S2.** Primers used in this study.

| **Name** | **Sequence (5’ to 3’)** | **Description** |
| --- | --- | --- |
| Construction of Gfp-reporters ^a^ | | |
| IVCE572 | **CATGTGGAAGACAGTGCC**ACCGGTTTCGAATTGACA | fwd - PconII (BpiI) |
| IVCE1120 | **TCACTGGAAGACAGTGCC**AAATATTCTGAAATGAGCTGTTGA | fwd - Ptrc (BpiI) |
| IVCE637 | **CATGTGGAAGACAGACG**CATCTTGTTGATACCCCCT | rev - riboswitch [B] (BpiI) |
| IVCE576 | **TCTCCTGAAGACGTACG**CATCTTGTTGTCCCTTGGT | rev - riboswitch C (BpiI) |
| IVCE578 | **TCACGTGAAGACAGACG**CATCTTGTTGCCTCCTTAGC | rev - riboswitch E* (BpiI) |
| IVCE574 | **CATGTGGAAGACCAG**CGTAAAGGAGAAGAACTTTTCA | fwd - Gfp-B0015 (BpiI) |
| IVCE575 | **CATGTGGAAGACCATCCC**TATAAACGCAGAAAGGCCC | rev - Gfp-B0015 (BpiI) |
| Domestication of BsaI site in pPMQAK1-T ^b^ | | |
| IVCE721 | CGCGAGATCCACGCTCACCGGCTCC | fwd - mutating BsaI-site in pPMQAK1 (in Amp^r^) |
| IVCE722 | AGCGTGGATCTCGCGGTATCATTGCAGCAC | rev - mutating BsaI-site in pPMQAK1 (in Amp^r^) |
| Domestication of BpiI sites in *cas9* ^a, b^ | | |
| IVCE711 | **ACGCAAGAAGACAGA**GGTTTCTTAGACGTCAGGT | fwd - pMD19 (BpiI) |
| IVCE712 | **ACGCAAGAAGACAGGT**TCCACACAACATACGAGC | rev - pMD19 (BpiI) |
| IVCE713 | **ACGCAAGAAGACCAGA**ACCGGTTTCGAATTGACA | fwd - PconII-Cas9 Piece 1 (BpiI) |
| IVCE714 | **ACGCAAGAAGACCACTT**ATCTTCTTCCACCAAAAAAGAC | rev - PconII-Cas9 Piece 1 (BpiI) |
| IVCE715 | **ACGCAAGAAGACCAT**AAGAAGCATGAACGTCATC | fwd - Cas9 Piece 2 (BpiI) |
| IVCE716 | **ACGCAAGAAGACCAT**TGCCTTCTCAAAATAGCATG | rev - Cas9 Piece 2 (BpiI) |
| IVCE717 | **ACGCAAGAAGACCTGCA**AGAGGACTTTTATCCATTTTTAAAAG | fwd - Cas9 Piece 3 (BpiI) |
| IVCE718 | **ACGCAAGAAGACCTT**ATCTTCTTTAAATGTCAAACTATCATC | rev - Cas9 Piece 3 (BpiI) |
| IVCE719 | **ACGCAAGAAGACTGGAT**ATTCAAAAAGCACAAGTGTCT | fwd - Cas9 Piece 4 (BpiI) |
| IVCE720 | **ACGCAAGAAGACTGACC**TTAGTCACCTCCTAGCTGA | rev - Cas9 Piece 4 (BpiI) |
| Construction of pPMQAK1-CRISPR/Cas9 base vectors ^a, c^ | | |
| IVCE1120 | **TCACTGGAAGACAGTGCC**AAATATTCTGAAATGAGCTGTTGA | fwd - Ptrc (BpiI) |
| IVCE1121 | **TCACTGGAAGACAGC**CATCTTGTTGATACCCCCT | rev - Ptrc + constant region + [B] + ATG (BpiI) |
| IVCE1128 | **GGATTGGAAGACCTC**CATCTTGTTGTCCCTTGGT | rev - Ptrc + constant region + [C] + ATG (BpiI) |
| IVCE1129 | **GCCATAGAAGACCTC**CATCTTGTTGCCTCCTTAGC | rev - Ptrc + constant region + [E*] + ATG (BpiI) |
| IVCE1122 | **TCACTGGAAGACAGATG**GATAAGAAATACTCAATAGGCTTAGA | fwd - Cas9 (no start) (BpiI) |
| IVCE1123 | **TCACTGGAAGACAGGTTA**TTAGTCACCTCCTAGCTGA | rev - Cas9 (extra stop) (BpiI) |
| IVCE1124 | **TCACTGGAAGACGTTAA**CCAGGCATCAAATAAAACGA | fwd - B0015 (BpiI) |
| IVCE1125 | **TCACTGGAAGACGTACCTCTAGTA**TATAAACGCAGAAAGGCCCAC | rev - B0015 + generic extra bp (BpiI) |
| IVCE1126 | **TCACTGGAAGACAGAG**GTTTGGAGACCACGTGTTCACAGCTTGTCTGTAAGCG | fwd - lacZ + (BsaI site for future use) (BpiI) |
| IVCE1127 | **TCACTGGAAGACAGTCCC**TGTGCCACGAGACCACGTGTGCAGCTGGCACGACAGGTTT | rev - lacZ + (BsaI site for future use) (BpiI) |
| IVCE1132 | **GCGTTAGAAGACGATGCC**GATAAGAAATACTCAATAGGCTTAGA | fwd - Cas9 (no start), for promoterless & ATG-less control construct (BpiI) |
| Mutagenesis of pPMQAK1-CRISPR/Cas9 base vectors to obtain extra variants ^b^ | | |
| IVCE1150 | CGACGAAAACTACGCTTTAGTAGCTTAATAACCAGGCATCAAATAAAACGAAA | fwd - add LVA-tag to Cas9 |
| IVCE1084 | AGCGTAGTTTTCGTCGTTTGCAGCGTCACCTCCTAGCTGACTCA | rev - add LVA-tag to Cas9 |
| IVCE1151 | GCTCGTATTGTGTGTGGAATTGTGAGCGG | fwd - mutate Ptrc -10-box (to one in BBa_J23104: TATAAT->TATTGT) |
| IVCE1152 | CCACACACAATACGAGCCGGATGATTAATTGTC | rev - mutate Ptrc -10-box (to one in BBa_J23104: TATAAT->TATTGT) |
| IVCE1153 | GGCTCGGACTATGTGTGGAATTGTGAGCGG | fwd - mutate Ptrc -10-box (to one in BBa_J23116: TATAAT->GACTAT) |
| IVCE1154 | CACACATAGTCCGAGCCGGATGATTAATTGTCA | rev - mutate Ptrc -10-box (to one in BBa_J23116: TATAAT->GACTAT) |
| IVCE1155 | GGCTCGGATTGTGTGTGGAATTGTGAGCGG | fwd - mutate Ptrc -10-box (to one in BBa_J23117: TATAAT->GATTGT) |
| IVCE1156 | CACACACAATCCGAGCCGGATGATTAATTGTCA | rev - mutate Ptrc -10-box (to one in BBa_J23117: TATAAT->GATTGT) |
| Construction of donor DNA pieces ^a^ | | |
| IVCE1144 | **AACGTCGGTCTCGTTCG**GCGATGTAAACGGCCACAAA | fwd - H1 (yfp ∆20 bp) (BsaI) |
| IVCE1145 | **TATGGCGGATCTTG**CTTCTGCTTATCGGCCATAA | rev - H1 (yfp ∆20 bp) (overhang overlaps with H2) |
| IVCE1146 | **CCGATAAGCAGAAG**CAAGATCCGCCATAACATCG | fwd - H2 (yfp ∆20 bp) (overhang overlaps with H1) |
| IVCE1147 | **ACACGTGGTCTCGGCCA**AGTAGAGAGCGTTCACCGAC | rev - H2 (yfp ∆20 bp) (BsaI) |
| IVCE1192 | **AACGTCGGTCTCGTTCG**AGGGTGTTGGCCAAGGTA | fwd - H1 (yfp ∆2240 bp) (BsaI) |
| IVCE1193 | **GCATCACCGGAAGCTAAA**TGGTTTCTCAGATTGCAGTTG | rev - H1 (yfp ∆2240 bp) (overhang overlaps with H2) |
| IVCE1194 | **CTGCAATCTGAGAAACCA**TTTAGCTTCCGGTGATGCCC | fwd - H2 (yfp ∆2240 bp) (overhang overlaps with H1) |
| IVCE1195 | **ACACGTGGTCTCGGCCA**GATTTTGTTCAGCTAGTAACTG | rev - H2 (yfp ∆2240 bp) (BsaI) |
| IVCE1326 | **AACGTCGGTCTCGTTCG**CCCGTAGCTTCCGGTGGTAT | fwd - H1 (rbcL+FLAG) (BsaI) |
| IVCE646 | **CATTGGCTTACTTATCGTCATCATCCTTGTAGTC**GAGGGTATCCATGGCCTCGA | rev - H1 (rbcL+FLAG) (overhang overlaps with H2) |
| IVCE647 | **TCGACTACAAGGATGATGACGATAAGTAAGCCAA**TGTTTGGATTGTCGGAGTTGT | fwd - H2 (rbcL+FLAG) (overhang overlaps with H1) |
| IVCE1327 | **ACACGTGGTCTCGGCCA**TTCTTTATTTTCATCCAGGAGTTCC | rev - H2 (rbcL+FLAG) (BsaI) |
| IVCE1212 | **AACGTCGGTCTCGTTCG**GGGACGGACAGTTATCCTAA | fwd - H1 (NS1: Yfp-insertion and ∆30 bp) (BsaI) |
| IVCE1328 | **TACCGAGGTCTCGTT**AGGAGACTTTGGTGGGCT | rev - H1 (NS1: Yfp-insertion) (BsaI) |
| IVCE1329 | **TACCGAGGTCTCCCT**AACTGACTGACCACTGAC | fwd - PpsbA2-Yfp-B0015 (for insertion into NS1 (BsaI) |
| IVCE1330 | **TACCGAGGTCTCCGAT**TATAAACGCAGAAAGGCCC | rev - PpsbA2-Yfp-B0015 (for insertion into NS1 (BsaI) |
| IVCE1331 | **TACCGAGGTCTCCA**ATCCCTTCAGTGGTACTCC | fwd - H2 (NS1: Yfp-insertion) (BsaI) |
| IVCE1215 | **ACACGTGGTCTCGGCCA**ACCTACCTGTCCTGGGTTGA | rev - H2 (NS1: Yfp-insertion and ∆30 bp) (BsaI) |
| IVCE1213 | **ACCACTGAAGGGAT**AGGAGACTTTGGTGGGCTG | rev - H1 (NS1 ∆30 bp) (overhang overlaps with H2) |
| IVCE1214 | **CACCAAAGTCTCCT**ATCCCTTCAGTGGTACTCC | fwd - H2 (NS1 ∆30 bp) (overhang overlaps with H1) |
| IVCE1216 | **AACGTCGGTCTCGTTCG**ATTGATGGCATTCGGGAGCC | fwd - H1 (NS2 ∆30 bp) (BsaI) |
| IVCE1217 | **TATGTTCCGCTTGC**ATGCCTAGGGGTAAACCATC | rev - H1 (NS2 ∆30 bp) (overhang overlaps with H2) |
| IVCE1218 | **TTACCCCTAGGCAT**GCAAGCGGAACATAACGTAT | fwd - H2 (NS2 ∆30 bp) (overhang overlaps with H1) |
| IVCE1219 | **ACACGTGGTCTCGGCCA**TGAAGTTAAAACCGTTGAGG | rev - H2 (NS2 ∆30 bp) (BsaI) |
| IVCE1266 | **AACGTCGGTCTCGTTCG**AAAGTTGGGCACAACCATTT | fwd - H1 (NS3 ∆30 bp) (BsaI) |
| IVCE1267 | **GTAAACTCCGCAGGG**CTTTATTTTGTTCCCGGTAC | rev - H1 (NS3 ∆30 bp) (overhang overlaps with H2) |
| IVCE1268 | **CCGGGAACAAAATAAAG**CCCTGCGGAGTTTACATAATC | fwd - H2 (NS3 ∆30 bp) (overhang overlaps with H1) |
| IVCE1269 | **ACACGTGGTCTCGGCCA**CAGCTATAGTTACGATCTTG | rev - H2 (NS3 ∆30 bp) (BsaI) |
| IVCE1224 | **AACGTCGGTCTCGTTCG**CTATCCTATGCATGTCATGT | fwd - H1 (NS4 ∆30 bp) (BsaI) |
| IVCE1225 | **CGTACTCGCTTAAA**TCCTTGGTCTGTTCAACTAA | rev - H1 (NS4 ∆30 bp) (overhang overlaps with H2) |
| IVCE1226 | **GAACAGACCAAGGA**TTTAAGCGAGTACGCCCGAA | fwd - H2 (NS4 ∆30 bp) (overhang overlaps with H1) |
| IVCE1227 | **ACACGTGGTCTCGGCCA**CATAGATATTTGCCCAGTTA | rev - H2 (NS4 ∆30 bp) (BsaI) |
| IVCE1315 | **GGTTACGGTCTCCAG**ACCTACCTGTCCTGGGTT | rev - H2 NS1 (∆30 bp), multi: compatible with donor NS4 (∆30 bp) (BsaI) |
| IVCE1316 | **GGTTACGGTCTCCGT**CTATCCTATGCATGTCATGTG | fwd - H1 NS4 (∆30 bp), multi: compatible with donor NS1 (∆30 bp) (BsaI) |
| IVCE1313 | **GTTTCGGGTCTCGTT**ACCTACCTGTCCTGGGTT | rev - H2 (NS1 ∆30 bp), multi: compatible with donor NS3 (∆30 bp) (BsaI) |
| IVCE1314 | **GTTTCGGGTCTCCGT**AAAGTTGGGCACAACCAT | fwd - H1 (NS3 ∆30 bp), multi: compatible with donor NS1 (∆30 bp) (BsaI) |
| IVCE1277 | **GCATCTGGTCTCGAG**CAGCTATAGTTACGATCTTG | rev – H2 (NS3 ∆30 bp), multi: compatible with donor NS4 (∆30 bp) (BsaI) |
| IVCE1241 | **GCATCTGGTCTCGTG**CTATCCTATGCATGTCATGTG | fwd – H1 (NS4 ∆30 bp), multi: compatible with donor NS3 (∆30 bp) (BsaI) |
| Construction of sgRNA pieces ^a, d^ | | |
| IVCE992 | **ACACGTGGTCTCGGTTT**TTGACAGCTAGCTCAGTCCT | fwd – BBa_J23117-sgRNA, to amplify whole sgRNA (BsaI) |
| IVCE993 | **AACGTCGGTCTCGCGAA**AAAAAAAGCACCGACTCGGTGC | rev – sgRNA scaffold, to amplify whole sgRNA (BsaI) |
| IVCE1142 | CTATGGCCGATAAGCAGAAGAATGTTTTAGAGCTAGAAATAGCAAGT | fwd - Cas9-handle (overhang: yfp-spacer) |
| IVCE1143 | AACATTCTTCTGCTTATCGGCCATAGCTAGCACAATCCCTAGG | rev – BBa_J23117 (overhang: yfp-spacer) |
| IVCE645 | CTAGGCCATGGATACCCTCTAAACGTTTTAGAGCTAGAAATAGCAAGT | fwd - Cas9-handle (overhang: rbcL-spacer) |
| IVCE644 | AACGTTTAGAGGGTATCCATGGCCTAGCTAGCACAATCCCTAGGACT | rev – BBa_J23117 (overhang: rbcL-spacer) |
| IVCE1204 | CTAGCCCACCAAAGTCTCCTATGGTTTTAGAGCTAGAAATAGCAAGT | fwd - Cas9-handle (overhang: NS1-spacer) |
| IVCE1205 | AACCATAGGAGACTTTGGTGGGCTAGCTAGCACAATCCCTAGG | rev – BBa_J23117 (overhang: NS1-spacer) |
| IVCE1206 | CTATGGTTTACCCCTAGGCATCAGGTTTTAGAGCTAGAAATAGCAAGT | fwd - Cas9-handle (overhang: NS2-spacer) |
| IVCE1207 | AACCTGATGCCTAGGGGTAAACCATAGCTAGCACAATCCCTAGG | rev – BBa_J23117 (overhang: NS2-spacer) |
| IVCE1262 | CTACCGGGAACAAAATAAAGTGCGTTTTAGAGCTAGAAATAGCAAGT | fwd - Cas9-handle (overhang: NS3-spacer) |
| IVCE1263 | AACGCACTTTATTTTGTTCCCGGTAGCTAGCACAATCCCTAGG | rev – BBa_J23117 (overhang: NS3-spacer) |
| IVCE1210 | CTAGTTGAACAGACCAAGGAACAGTTTTAGAGCTAGAAATAGCAAGT | fwd - Cas9-handle (overhang: NS4-spacer) |
| IVCE1211 | AACTGTTCCTTGGTCTGTTCAACTAGCTAGCACAATCCCTAGG | rev – BBa_J23117 (overhang: NS4-spacer) |
| Constructing sgRNA-array template vectors as described in Li et al.^2^ ^e^ | | |
| IVCE851 | **CTAGAATTCGCGGCCGCTTCTAGA**ACACGTGGTCTCGGTTTTTGACAGCTAGCTCAGTC | fwd – BBa_J23117-sgRNA, adds BsaI (EcoRI+NotI+XbaI) |
| IVCE852 | **CTGCAGCGGCCGCTACTAGT**AACGTCGGTCTCGCGAAAAAAAAAGCACCGACTCG | rev – sgRNA scaffold, adds BsaI (PstI+NotI+BcuI) |
| IVCE853 | **CTAGAATTCGCGGCCGCTTCTAGA**GTTTTAGAGCTAGAAATAGCAAG | fwd - Cas9-handle (EcoRI+NotI+XbaI) |
| IVCE854 | **CTGCAGCGGCCGCTACTAGT**TAGCTAGCACAATCCCTAGGACTGAGCTAGCTGTCAAAAAAAAAGCACCGACTCGGTGC | rev – sgRNA scaffold, adds BBa_J23117 overhang (PstI+NotI+BcuI) |
| For sgRNA-array construction (multiplex targeting), as described in Li et al.^2^ ^a, d^ | | |
| IVCE1228 | **GGACTAGAAGACGAT**AGTTGAACAGACCAAGGAACAGTTTTAGAGCTAGAAATAGCAAGT | fwd - Cas9-handle (in pMD19 template vector), adds NS4-spacer (BpiI) |
| IVCE1229 | **GGACTAGAAGACGAGGC**TAGCTAGCACAATCCCTAGG | rev – BBa_J23117 (in pMD19 template vector), overhang compatible with NS1-spacer (BpiI) |
| IVCE1230 | **GGACTAGAAGACTG**AGCCCACCAAAGTCTCCTATGGTTTTAGAGCTAGAAATAGCAAGT | fwd - Cas9-handle, adds NS1-spacer (BpiI) |
| IVCE1235 | **GGACTAGAAGACGAAC**TAGCTAGCACAATCCCTAGG | rev – BBa_J23117, overhang compatible with NS4-spacer (BpiI) |
| IVCE1273 | **TAACCCGAAGACAGGG**TAGCTAGCACAATCCCTAGG | rev – BBa_J23117, overhang compatible with NS3-spacer (BpiI) |
| IVCE1274 | **TAACCCGAAGACTCT**ACCGGGAACAAAATAAAGTGCGTTTTAGAGCTAGAAATAGCAAGT | fwd - Cas9-handle, adds NS3-spacer (BpiI) |
| Construction of P*nrsD*-Cure-sgRNA pieces | | |
| IVCE1257 | **TTACGGGAAGACTGCAT**TATTCGATTCAGTACCAAGTAC | fwd - PnrsD (BpiI) |
| IVCE1258 | AACCGATCCCCGGGAAAACAGCATAATTGATCTCTATTCTATGCCC | rev - PnrsD (overhang: Cure-sgRNA nr 1 spacer) |
| IVCE1259 | AACAAGCGGCTAAAAGCGCTCCAGCGTAATTGATCTCTATTCTATGCCC | rev - PnrsD (overhang: Cure-sgRNA nr 2 spacer) |
| IVCE1260 | AACAGCAAAAGTTCGATTTATTAATTGATCTCTATTCTATGCCC | rev - PnrsD (overhang: Cure-sgRNA nr 3 spacer) |
| IVCE1261 | AACAAAATCTCTGATGTTACATTAATTGATCTCTATTCTATGCCC | rev - PnrsD (overhang: Cure-sgRNA nr 4 spacer) |
| IVCE1178 | **TTACGGGAAGACTGTCCC**TGATAACGGACTAGCCTTATT | rev – sgRNA scaffold (+54 truncation) (BpiI) |
| IVCE1180 | CTATGCTGTTTTCCCGGGGATCGGTTTTAGAGCTAGAAATAGCAAGT | fwd - Cas9-handle (overhang: Cure-sgRNA nr 1 spacer) |
| IVCE1182 | CTACGCTGGAGCGCTTTTAGCCGCTTGTTTTAGAGCTAGAAATAGCAAGT | fwd - Cas9-handle (overhang: Cure-sgRNA nr 2 spacer) |
| IVCE1184 | CTAATAAATCGAACTTTTGCTGTTTTAGAGCTAGAAATAGCAAGT | fwd - Cas9-handle (overhang: Cure-sgRNA nr 3 spacer) |
| IVCE1186 | CTAATGTAACATCAGAGATTTTGTTTTAGAGCTAGAAATAGCAAGT | fwd - Cas9-handle (overhang: Cure-sgRNA nr 4 spacer) |
| Colony-PCR screening primers | | |
| IVCE799 | ATGGTATCCAAAGGCGAGGAG | fwd - yfp ∆20 bp |
| IVCE1170 | TTGCACACTTCCGTCTTCGA | rev - yfp ∆20 bp |
| IVCE141 | TCGATGGAGGGTCAGACCAT | fwd - yfp ∆2240 bp |
| IVCE142 | TCAGCAAAACTGCCGCAATC | rev - yfp ∆2240 bp |
| IVCE663 | GCCGTAGTTGACCGTCAGAA | fwd - rbcL-FLAG |
| IVCE662 | GCAGACTCAACCCCGAAGAA | rev - rbcL-FLAG |
| IVCE1278 | ATTTACACCAGCGCCGGTTT | fwd - NS1 (Yfp and ∆30 bp) |
| IVCE1279 | ACCGCTAAACCCACCTCTTG | rev - NS1 (Yfp and ∆30 bp) |
| IVCE1280 | GCTGGTTTGGGGTGTTGATG | fwd - NS2 ∆30 bp |
| IVCE1281 | ACCCCAGCTACCCCTAACAT | rev - NS2 ∆30 bp |
| IVCE1323 | TGGTGGCTAAGTTGTACCGG | fwd - NS3 ∆30 bp |
| IVCE1324 | AGTAACTTACGACGGGTGGG | rev - NS3 ∆30 bp |
| IVCE1284 | TACCAAGAACTACTGCGGCG | fwd - NS4 ∆30 bp |
| IVCE1285 | CTGGGTCGATCTTCCTTCCA | rev - NS4 ∆30 bp |
| IVCE174 | GGAGAAAACTCACCGAGGCA | fwd - Km^r^ screening (curing) |
| IVCE175 | CTCAGGCGCAATCACGAATG | rev - Km^r^ screening (curing) |

^a^ Golden Gate cloning compatible overhangs are in bold. See description column for specification of the type II restriction enzyme in question.

^b^ Mutated nucleotide indicated in red.

^c^ Underlined sequences shows the BsaI-containing extra region added to the CRISPR/Cas9 base vectors, for use when constructing the target vector.

^d^ Underlined sequences are the added sgRNA spacers.

^e^ Overhangs with BioBrick cloning sites are indicated in bold, restriction sites are specified in the description column. Underlined indicates the added BsaI-containing regions added to flank the finished sgRNA-array. In IVCE854 the BBa_J23117 promoter added after the sgRNA scaffold (Cas9-handle and S. pyogenes terminator) is also underlined.

**Table S3.** Details for all tested sgRNA-spacers.

| Target | Spacer sequence (5’-3’) | PAM | On-target  score ^a^ |
| --- | --- | --- | --- |
| *yfp* | ATGGCCGATAAGCAGAAGAAT | GGG | 45.8 |
| *rbcL* | AGGCCATGGATACCCTCTAAAC | CGG | 44.2 |
| NS1 (*slr0168*) | AGCCCACCAAAGTCTCCTATG | TGG | 62.5 |
| NS2 (*slr1181*) | ATGGTTTACCCCTAGGCATCAG | TGG | 62.0 |
| NS3 (*slr2030-2031*) | ACCGGGAACAAAATAAAGTGC | AGG | 57.5 |
| NS4 (*slr0397*) | AGTTGAACAGACCAAGGAACA | GGG | 73.0 |
| pPMQAK1, nr 1 ^b^ | ATGCTGTTTTCCCGGGGATCG | CAG | 1.9 |
| pPMQAK1, nr 2 ^b^ | ACGCTGGAGCGCTTTTAGCCGCTT | TAG | 5.7 |
| pPMQAK1, nr 3 ^b^ | AATAAATCGAACTTTTGCT | GAG | 39.2 |
| pPMQAK1, nr 4 ^b^ | AATGTAACATCAGAGATTTT | GAG | 7.9 |

^a^ The on-target score as determined by Benchling.^3^ This score is determined by using an algorithm based on mammalian cell data, it also doesn’t consider the position of the spacer in the target gene.^4^ However, note that this scoring might not be directly applicable for a prokaryotic host.^5^

^b^ The pPMQAK1-backbone targeting spacers used in the Cure-sgRNAs (nr 1-4).

**Table S4.** Off-target analysis of sgRNAs used in this study, against the S6803 genome, using the CasOT software.^6^ The PAM was specified to NGG only.

|  | Target | | | | | | Cure-sgRNA | | | |
| --- | --- | --- | --- | --- | --- | --- | --- | --- | --- | --- |
| Type ^a^ | *yfp* | *rbcL* | NS1 | NS2 | NS3 | NS4 | 1 | 2 | 3 | 4 |
| A06 |  |  |  |  |  |  |  |  |  | 1 |
| A13 |  |  |  |  |  |  |  |  | 1 |  |
| A14 |  |  |  |  |  | 1 | 2 |  | 5 | 2 |
| A15 |  |  |  |  |  | 1 |  |  |  | 1 |
| A16 | 1 |  |  | 1 | 1 | 1 |  | 1 |  |  |
| A17 |  |  |  |  | 1 | 1 |  |  |  |  |
| A18 | 4 | 1 |  |  |  |  |  | 1 |  |  |
| A19 | 1 |  |  |  |  |  |  | 1 |  |  |
| A110 |  |  |  |  |  |  |  | 1 |  |  |
| A23 | 1 | 1 |  |  |  |  |  |  | 6 | 1 |
| A24 | 4 | 1 | 2 |  | 2 | 2 | 3 |  | 17 | 2 |
| A25 | 8 |  | 2 | 3 | 6 | 8 | 8 |  | 16 | 4 |
| A26 | 10 | 1 | 4 | 3 | 10 | 9 | 63 | 1 | 12 | 8 |
| A27 | 22 | 4 | 3 | 9 | 14 | 8 | 14 | 3 | 7 | 3 |
| A28 | 19 | 6 | 6 | 2 | 11 | 5 | 6 | 15 |  |  |
| A29 | 7 | 3 | 1 | 2 | 7 |  |  | 13 |  |  |
| A210 |  |  |  |  |  |  |  | 18 |  |  |
| A211 |  |  |  |  |  |  |  | 6 |  |  |
| A212 |  |  |  |  |  |  |  | 2 |  |  |

^a^ Indicates the type of mismatch. The first number (0-2) specifies the number of mismatches in the seed region (12 nt proximal to PAM) of the binding sequence. The following number indicates the mismatches in the remaining non-seed region.

**Sequences**

**P*_conII_*:** (TSS in bold)

ACCGGTTTCGAATTGACAATTAATCATCGGCTCGTATAATGGTAC**C**

**P*_trc_*:** (TSS in bold, constant region as in Nakahira et al.^7^ is underlined)**:**

AAATATTCTGAAATGAGCTGTTGACAATTAATCATCCGGCTCGTATAATGTGTGG**A**ATTGTGAGCGGATAACAATTTCATACGCTCACAATTGGTACC

**Riboswitch B:** (as in Topp et al.^8^, aptamer in bold, start codon underlined)

**GGTGATACCAGCATCGTCTTGATGCCCTTGGCAGCACC**CGCTGCGCAGGGGGTATCAACAAGATG

**Riboswitch C:** (as in Topp et al.^8^, aptamer in bold, start codon underlined)

TGATAAGATAGG**GGTGATACCAGCATCGTCTTGATGCCCTTGGCAGCACC**AAGGGACAACAAGATG

**Riboswitch E*:** (as in Topp et al.^8^, aptamer in bold, start codon underlined)

**GGTGATACCAGCATCGTCTTGATGCCCTTGGCAGCACC**CTGCTAAGGAGGCAACAAGATG

**P*_nrsD_*-Cure-sgRNA nr 4:**

P*_nrsD_* in blue, its two mapped TSS^9^ in bold; the Cure-sgRNA nr 4 spacer is underlined, the Cas9-handle is in green, and yellow shows the +54 truncated *S. pyogenes* terminator as described in Hsu et al.^10^.

TATTCGATTCAGTACCAAGTACTATTGCGGGGACAGGACGTTTCTCAAGGCCCTCATCAATATCCCCCCTGGGGGCATAGAATAGAGAT**C**AAT**T**AATGTAACATCAGAGATTTTGTTTTAGAGCTAGAAATAGCAAGTTAAAATAAGGCTAGTCCGTTATCA

**Supporting methods - Using the inducible CRISPR/Cas9-system**

For an overview of the CRISPR/Cas9 target vector construction and subsequent workflow in S6803, see Figure 1 in the main text.

**Theophylline inducer stock: preparation and use**

A 200 mM theophylline stock, dissolved in 100% DMSO, is used. This does not dissolve at RT and requires heating before use. Theophylline is very stable^11^ and repeated heating was not found to reduce the stocks performance. To dissolve it, heat in a water bath at 42-50°C, mix a few times by vortexing. When fully dissolved, use directly to make inducer-supplemented BG11-plates (a final concentration of 0.25 mM is recommended), or for other applications. Store at 4°C between uses.

**Available pPMQAK1-CRISPR/Cas9 base vectors**

The best performing pPMQAK1-CRISPR/Cas9 base vectors described in this study will be submitted to Addgene. These are the constructs in order of expression strength: [E*](104) > [B]. The [E*](104) and [B] constructs are also available supplemented with P*_nrsD_*-Cure-sgRNA nr 4.

**Designing and constructing sgRNAs**

Identification of suitable protospacers in the target area can be done with e.g. Benchling.^3^ Useful guidelines are to select spacers that target the template strand,^12^ that guide Cas9 to cut as close to the edit site as possible,^13^ that don’t have significant off-target binding, and that doesn’t have extreme GC-content (>75%, <25%). In this study, spacers were also chosen to have an A at the TSS (often by adjusting spacer length by a few nts), however it is unknown if this is strictly necessary for P_BBa_J23117_.

An sgRNA sequence example:

TTGACAGCTAGCTCAGTCCTAGGGATTGTGCTAGCT**A**TGGCCGATAAGCAGAAGAATGTTTTAGAGCTAGAAATAGCAAGTTAAAATAAGGCTAGTCCGTTATCAACTTGAAAAAGTGGCACCGAGTCGGTGCTTTTTTT

(Blue: P_BBa_J23117_, bold: TSS, underlined: spacer, green: Cas9-handle, yellow: *S. pyogenes* terminator).

To make the sgRNA piece compatible for BsaI-based Golden Gate cloning, the following overhangs are required on the 5’- and 3’-ends of the above sgRNA. (BsaI-site shown in bold, created overhangs upon digestion are underlined):

- Fwd primer overhang: 5’-ACACGT**GGTCTC**GGTTT-….
- Rev primer overhang: 5’-AACGTC**GGTCTC**GCGAA-….

For a multiple-target construct

When multiplexing, an sgRNA-array needs to be constructed; this sgRNA-array must have the same BsaI-containing overhangs as described above. In this study this was done using the method described by Li et al.^2^ The following template vectors built in this study were used for this purpose: pMD19-BsaI- P_BBa_J23117_-sgRNA-BsaI-Sp^r^, and pMD19-Cas9_handle-*S.pyogenes*_terminator- P_BBa_J23117_-Cm^r^.

**Designing and constructing Donor DNAs**

In this study, homology arms of roughly 350 bp were used on either side of the intended edit. However, for e.g. large insertions it could be beneficial to use longer homology arms.^2^

Donor DNAs are preferably constructed by overlap-PCR to join the two homology arms together to one piece. To create the desired edit, the primers used to amplify the homology regions are designed accordingly. The donor DNA must also be designed to mutate or remove the PAM and preferably also parts of the proximal seed-sequence of the protospacer. See the study by Jiang et al. for advice on which mutations are effective.^14^ For larger insertions the donor DNA segments can be added as separate pieces to the Golden Gate assembly.

To make the donor DNA piece compatible for BsaI-based Golden Gate cloning, the following overhangs are required on the outermost 5’- and 3’-ends. (BsaI-site shown in bold, created overhangs upon digestion are underlined):

- Fwd primer overhang: 5’-AACGTC**GGTCTC**GTTCG-….
- Rev primer overhang: 5’-ACACGT**GGTCTC**GGCCA-….

For a multiple-target construct

For multiplexed targeting the above overhangs must be used for the forward primer of the first donor DNA, and the reverse primer for the last donor DNA, respectively. The rest of the primers must be designed (e.g. with Benchling)^3^ to have BsaI-overhangs that allow for assembly of the different donor-DNAs in the desired order.

**Golden Gate assembly of the pPMQAK1-CRISPR/Cas9 target vector**

All parts (sgRNA, donor DNA, and pPMQAK1-CRISPR/Cas9 base vector) are mixed with BsaI and T4 DNA ligase in a one-pot reaction. A 10 µl reaction is often enough; the volume can be increased to accommodate for e.g. assembly of many fragments for a multiplexing target vector.

- 20 ng pPMQAK1-CRISPR/Cas9 base vector. (No prior digestion is needed!)
- Add a 2:1 molar ratio of each insert (sgRNA, donor DNA).
- Add water so that the total final reaction volume reaches 10 µl (or as decided).
- Add the T4 DNA ligase buffer.
- Add the BsaI and T4 DNA ligase, usually 0.5-1 µl each per reaction.
- Mix and place in the thermocycler (see below).

In the thermocycler, 30 cycles are noted below. However, fewer cycles (e.g. 10) will also work but yield more background colonies.

1. 37°C – 5 min (digestion)
2. 16°C – 10 min (ligation)
3. Cycle back to step 1+2, x30
4. 55°C – 5 min (enzyme inactivation)
5. 10°C ∞

Transform your *E. coli* using 2-5 µl of the Golden Gate reaction. The pPMQAK1-CRISPR/Cas9 vectors are blue/white-screening compatible. For a sufficiently concentrated plasmid preparation (70-100 ng/µl), miniprep 10 ml *E. coli* culture. Confirm that the construct is correct by sequencing (covering the sgRNA and donor DNA).

**Transforming S6803 with the pPMQAK1-CRISPR/Cas9 target vector**

- Use 5-10 ml (OD_730_ 0.6-1) per electroporation.
- Pellet cells (10 min, 4500 rpm, 4°C), wash in cold sterile water. Repeat a total of three times. To reduce time, wash all cells together.
- Re-suspend washed cells in enough cold sterile water so that 50 µl can be aliquoted per electroporation into separate eppendorf tubes.
- Add up to 5 µl (100-350 ng DNA) of the vector. Incubate on ice for ~10 min before transferring to cooled electroporation cuvettes (2 mm gap).
- Electroporate using a Bio-Rad MicroPulser, setting EC2 (V=2.5 kV), or similar.
- Place cuvettes back on ice.
- Collect cell suspensions from the cuvettes and suspend in 3 ml BG11, recover for 16-24 h under standard growth conditions.
- Prepare selective (+Km) BG11 plates without and with 0.25 mM theophylline. If a P*_nrsD_*-Cure-sgRNA supplemented vector is used, use cobalt-free BG11.
- Collect recovered cells, pellet (10 min, 4500 rpm, 4°C) and re-suspend in ~500 µl BG11 (or cobalt-free BG11).
- Plate ~250 µl each on selective plates without and with inducer.
- Incubate under normal growth conditions. Colonies will start appearing after 1 week; allow them to grow a bit bigger for a total of about 10-14 days.

With a functional sgRNA and non-leaky CRISPR/Cas9, no or a few colonies will appear on the inducer-plate, while the no-inducer plate will have many colonies (see picture below). If any colonies appear on the inducer-plate, screen these for the desired edit.

**Inducing CRISPR/Cas9-mediated genome editing**

From the non-inducer transformation plates, pick healthy colonies but avoid the largest colonies due to the risk of these having mutated the CRISPR/Cas9 system. Avoid doing pre-cultures as this can enrich cells that have “escaped” the CRISPR/Cas9-system.

- Prepare selective BG11 plates with 0.25 mM theophylline. (If P*_nrsD_*-Cure-sgRNA vectors are used, use cobalt-free BG11).
- The colonies, preferably triplicates, are picked and suspended in ~80 µl BG11 (or cobalt-free BG11).
- Prepare dilutions, e.g. a three-step 5x dilution-series, which gives 5x, 25x, and 125x dilution suspensions. A final volume of 80-100 µl of each is enough.
- Plate the suspensions (see strategy below), ~30-50 µl suspension per section.
- To control that the induced colonies harbor an active CRISPR/Cas9-system, one can do a spot assay on plates with and without inducer. Equal survival on both plate-types indicates a likely “escaped” mutated colony.
- Incubate for around 10-14 days, until large green colonies are available.

Since the editing efficiency is unknown for new targets, 2-3 different dilutions of the suspended cells can be plated on inducer-plates (see picture below). If a difficult edit is anticipated (e.g. multiplexed editing or large insertion, see right plate below), plate the remaining no-dilution suspension and 5x dilution. If an easier edit is expected (e.g. a small deletion, see left plate below) one can instead plate the 5x-125x suspensions.

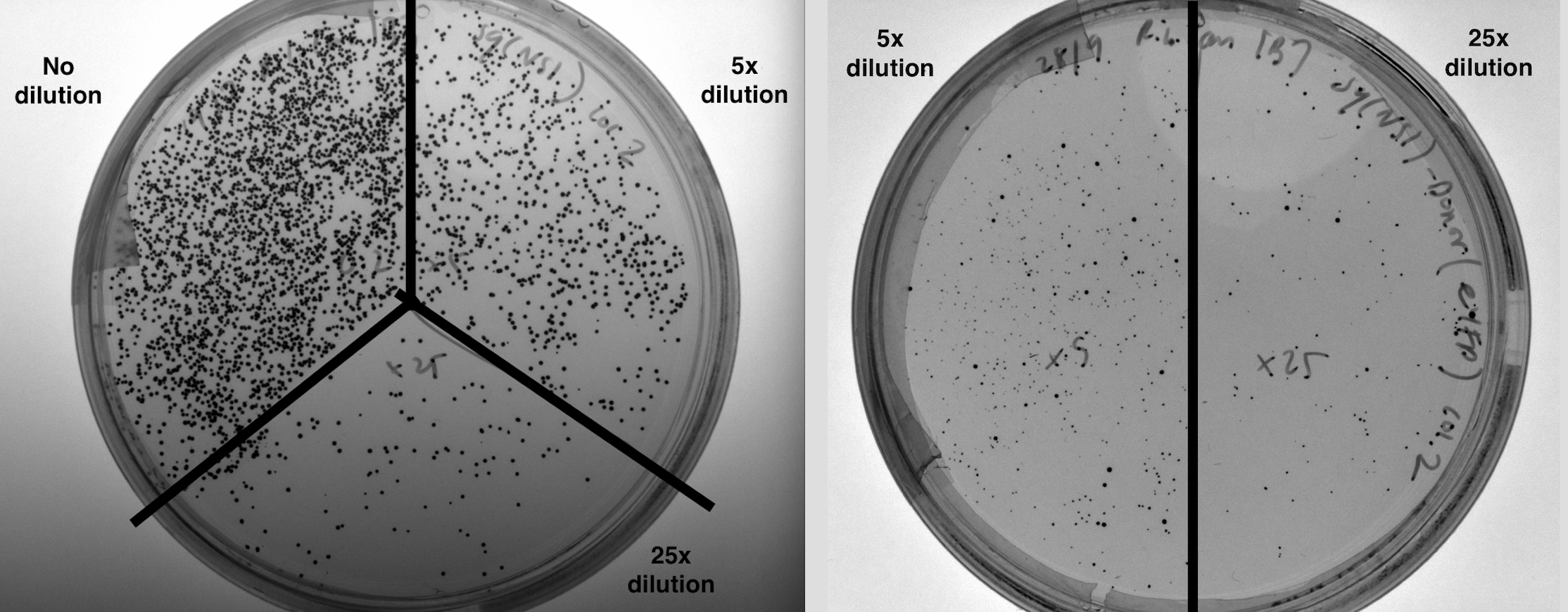

S**creening for genome editing**

For colony-PCR, ensure that at least one primer binds outside the homology regions used in the donor DNA to avoid false positives. Pick and screen colonies with a healthy phenotype. If multiple transformants were induced, screen colonies from inducer-plates where cells appear to not harbor an inactive CRISPR/Cas9-system (e.g. due to a cell-lawn, or consult a prepared spot assay). Any colonies with un-segregated edits can be streaked on new inducer-plates so that new single colonies will form.

Colonies picked for screening can preferably also be streaked on new non-inducer plates, so that any positives are readily available for the subsequent curing step.

**Curing pPMQAK1-CRISPR/Cas9 from edited cells**

The curing is performed in culture rather than on plates. Use regular BG11 here.

- Prepare BG11 with 2.5 µM Ni^2+^ and 0.25 mM theophylline. Add no Km!
- Pick fully edited colonies into the above media. Do small cultures (~2-3 ml).
- Grow for 4 days.
- Start a 2^nd^ cultivation round by diluting the first round 500x (still inducer-media).
- Grow the 2^nd^ round for 4 days. Plate on non-selective plates (see below).
- Incubate plates until colonies appear.
- Screen colonies for plasmid loss, e.g. by colony-PCR or streaking on Km-plates. Due to the low efficiency of the curing, start by screening 25 colonies.

To obtain separated colonies when plating:

- Measure OD_730_ of the 2^nd^ round cultures, dilute these to OD_730_ 0.2 in ~100 µl.
- Dilute this again, preferably by two 56x dilutions. A final ~100 µl is enough.
- Plate ~100 µl of the final dilution on non-selective, regular BG11 plates.
